# Effect of Age on Xenobiotic-Induced Autoimmunity

**DOI:** 10.1101/2025.05.22.655368

**Authors:** Caroline de Ocampo, Amy K. Peiss, Ho Yeung Leung, Lisa MF Janssen, Dwight H. Kono, Jessica M. Mayeux, K. Michael Pollard

**Author notes:** Corresponding author: K. M. Pollard, PhD, Department of Immunology and Microbial Science, MB120, The Scripps Research Institute, 10550 North Torrey Pines Road, La Jolla, CA 92037, USA.

## Abstract

Aging is associated with increased spontaneous autoantibody production and chronic inflammation, yet its impact on xenobiotic-induced autoimmunity remains unexplored. This study investigates the effect of age on mercury-induced autoimmunity (HgIA) in B10.S mice, a model of xenobiotic-induced autoimmunity characterized by anti-nucleolar autoantibodies (ANoA). Mature (3 months), adult (6 months), middle-aged (12 months), and old-age (24 months) mice were exposed to mercury (HgCl₂) or phosphate-buffered saline (PBS) for 4–5 weeks. While spontaneous anti-nuclear antibodies (ANA) increased with age in PBS-treated mice (34% in middle-aged, 57% in old age mice), HgIA incidence declined in old age mice, with only 59% (26/44) developing significant ANoA titers compared to 91–100% in younger cohorts. Notably, 56% (10/18) of initially ANoA-negative old mice had detectable ANoA at a lower dilution, indicating a reduced but not absent response. ANoA negativity in old age mice was associated with lower immunoglobulin levels, reduced anti-chromatin antibodies, and diminished germinal center formation, suggestive of immunosenescence. Flow cytometry revealed age-related declines in CD4⁺ T cells, with mercury exposure augmenting T-cell differentiation in younger but not old mice. These findings demonstrate that aging enhances spontaneous autoimmunity but impairs xenobiotic-induced autoimmunity, with a subset of old age mice retaining partial responsiveness at lower dilutions, highlighting the complex interplay between immunosenescence and environmental triggers.

## Introduction

Many environmental and xenobiotic exposures have been linked to autoimmunity in human populations [1–3]. For instance, silica dust is an established risk factor for systemic autoimmune diseases [1, 4], while mercury exposure can trigger pathological outcomes—such as autoantibodies, inflammatory markers, and renal pathology—without a definitive association with a diagnosable autoimmune disease [1, 5, 6]. Animal studies support these associations [2, 7–11]. Autoantibodies are a serological hallmark of autoimmune diseases, with anti-nuclear antibodies (ANA) being characteristic of systemic autoimmune diseases [12, 13] where they have been associated with disease manifestations and severity and play a critical role in the preclinical or subclinical stages of autoimmunity. In fact, diagnostically relevant autoantibodies and inflammatory markers appear in the bloodstream a decade or more before the clinical diagnosis of several autoimmune diseases [14–18]. Exposure to xenobiotics is also linked to systemic inflammation and autoantibodies that emerge prior to symptomatic autoimmune disease [1, 2, 19].

Aging is associated with chronic low-grade inflammation, termed *inflammaging* [20, 21], as well as a progressive decline in immune function (i.e., immune senescence or immuno-senescence) manifested by poor responses to vaccination, increased susceptibility to infection, and chronic diseases [22–26]. Research highlights chronic inflammation—driven by proinflammatory cytokines and the senescence-associated secretory phenotype (SASP)—as a predictor of deteriorating immune health [24]. Concomitant with these changes, aging is associated with a rise in subclinical autoimmunity, including autoantibody production [21, 23, 24, 27, 28]. A specific subset of B cells, called age-associated B cells (ABCs), which produce autoantibodies and accumulate with age in both humans and mice [29–31], arise earlier and more prominently in many autoimmune-prone mouse strains [30]. Whether these subclinical markers of autoimmunity— autoantibodies, inflammatory markers, and ABCs—signal the onset of inflammaging remains unclear. These observations suggest that autoreactive B cells are not only central to autoimmunity [32] but also play a role in the aging process [33]. Significantly, the interplay between aging and xenobiotic-induced autoimmunity remains largely unexplored, beyond the recognition that prolonged occupational exposure heightens risk [1].

Xenobiotic exposures can occur in many ways and thus be present at different stages of life. Similarly, autoimmune diseases can occur throughout life, although the onset of many occurs in early and mature adult life [34, 35]. While aging is associated with an increase in subclinical autoimmunity, particularly autoantibodies [27, 36] in humans [37, 38] and mice [39, 40], the incidence of autoimmune diseases does not increase in later life [24, 27]. This lack of increase, however, does not imply that older individuals are invulnerable to xenobiotic-induced autoimmunity, or that lifetime exposure to xenobiotics is unrelated to subclinical autoimmunity.

To study the relationship between age and xenobiotic-induced autoimmunity we exposed mature (3 months old), adult (6 months old), middle aged (12 months old) and old age (24 month old) B10.S mice, equivalent to the human life phases of 19-26 years, 30-35 years, 40-48 years, and 65-75 years old respectively, to mercury [41]. Mercury-induced autoimmunity (HgIA) is characterized by a highly-specific MHC class II restricted anti-nucleolar autoantibody (ANoA) response in susceptible mouse strains [42, 43]. ANoA is a type of ANA that can be distinguished by a staining pattern localized to the nucleolus. Antibodies to fibrillarin, a protein found within the nucleolus involved in ribosomal RNA processing, comprise the predominant ANoA response of HgIA. As observed with other mouse strains [39, 44–46] aging in B10.S mice was associated with the spontaneous development of ANA. However, our findings include the novel observation that the incidence of HgIA declines in old age. Moreover, 40% of old age mice did not produce ANoA upon mercury exposure, and failure to respond was associated with reduced immunoglobulins and germinal center formation.

## Materials and Methods

### Mice

Male and female B10.S/SgMcdJ (B10.S) mice, as described previously [47], were utilized for all experiments. All mice were bred and maintained under pathogen-free conditions at The Scripps Research Institute (TSRI) Animal Facility (La Jolla, CA). Animals were kept at 68-72°F and 60-70% humidity, with a 12/12h light-dark cycle and sterilized cages replaced every two weeks with fresh water and food (Global 18% Protein Rodent Diet (2018), Teklad, Envigo, Madison, WI, USA).

### Aging of mice

Cohorts of B10.S mice were bred and maintained until they reached 2 months for mature mice, 5 months for adult mice, 10 months for middle aged mice, and 22 months for old age mice, which approximates the following life phase equivalencies; Mature adult (Mice:3-6 mo, Human:20-30yr), middle age (Mice:10-14 mo, Human:38-47yr), and old age (Mice:18-24 mo, Human:56-69yr) [41]. Survival curves for several cohorts showed 75% survival at 21 months for females and 22 months for males which is similar to what has been previously established [48] (Supplemental Figure 1). Mice were included in the study on a rolling basis as they reached the appropriate age. Therefore, experimental data is representative of several individual cohorts pooled together. During aging and experimental manipulation, animals were observed daily for signs of debilitation or illness based on the TSRI Guidelines for Animals Used in Aging Research (Supplemental Table 3). Number of animals per group is expressed as (n= # of mice treated with PBS or HgCl_2_).

### Induction of autoimmunity

Male and female mice were injected subcutaneously with 40 µg HgCl_2_ (Mallinckrodt Baker, Phillipsburg, NJ) in 100 µl PBS twice per week for 4 to 5 weeks. Controls received 100 µl PBS alone. This HgCl_2_ dose is consistent with occupational exposure [49, 50]. Procedures using HgCl_2_ were approved by the TSRI Institutional Animal Care and Use Committee and the TSRI Department of Environmental Health and Safety.

### Serology

Anti-nuclear antibodies (ANAs) were determined by indirect immunofluorescence as described previously [47]. HEp-2 cells on glass slides (Inova Diagnostics Inc., San Diego, CA) were incubated with serum diluted 1:100 and then with 1:400 dilution of Alexa Fluor 488–conjugated goat anti-mouse IgG (Molecular Probes, Carlsbad, CA). Each incubation period was followed by washes with PBS. Slides were mounted with Vectashield Mounting Medium (Vectorlabs, Burlingame, CA) and observed using an Olympus BH2 microscope (Olympus Corp., Tokyo, Japan). Digital images of ANA patterns were captured using a LEICA DFC 365 FX camera and analyzed using Leica Application Suite AF software (Leica Microsystems, Buffalo Grove, IL, USA). ANAs were scored for intensity (0 to 4 scale) and pattern under blinded conditions. The presence of nucleolar staining was scored as a separate category to indicate the presence of anti-nucleolar autoantibodies (ANoA). ANA scores of 1 or greater were considered positive while for ANoA, scores of 0.5 or greater were considered positive. ELISAs were used to quantitate autoantibodies against chromatin, and ENA5 (containing Sm, RNP, Scl-70, SS-A and SS-B antigens) (Inova Diagnostics Inc.) as previously described [9], and IgG and IgG1 (Immunology Consultants Laboratory, Newburg, OR) according to the manufacturer’s instructions. Indirect ELISA was used to quantitate IgG2c using goat anti-mouse IgG2c (Southern Biotech, Birmingham, AL) capture antibody and HRP-conjugated goat anti-mouse IgG (Invitrogen, Carlsbad, CA) detection antibody. ELISAs were evaluated using a Spectra Max M3 spectrophotometer (Molecular Devices, San Jose, CA) at 450 nm using Softmax Pro 6.3 software (Molecular Devices). The sum of the mean plus two standard deviations for 3 month old (mature) PBS controls was used to discriminate between positive and negative responses.

### Flow cytometry

Single-cell suspensions were prepared from spleens by mechanical homogenization through a 70 µm strainer (Fisher Scientific, Hanover Park, IL), incubation with Red Blood Cell Lysing Buffer Hybri-Max™ (Sigma Life Science, Burlington, MA), and re-suspension in FACS buffer (PBS with 4% FBS). One million cells were incubated with TruStain FcX^TM^ (anti-mouse CD16/32) antibody (BioLegend, San Diego, CA) to block Fc receptors before cells were stained with 19 anti-mouse antibodies for specific cell surface markers (Supplemental Table 1) in BD Horizon Brilliant Stain Buffer (BD Biosciences). All incubation periods were followed by washes with FACS buffer. Data was acquired using a Cytek Aurora 5-Laser Flow Cytometer and SpectroFlo V3.2.1 (Cytek Biosciences, Fremont, CA) and analyzed with FlowJo V10.10.0 (Tree Star, Ashland, OR). Populations were determined by excluding debris, aggregates and dead (DAPI stained) cells, then separated into TCRb positive and negative populations. Subset populations were identified as described in the results and summarized in Supplemental table 2.

### Statistics

Data was analyzed using Graphpad Prism V10 (GraphPad Software, San Diego, CA) and is visualized as individual mice and group mean with SEM. Unpaired two-tailed Mann–Whitney U test was used for comparison between PBS and HgCl_2_ exposure within an age group. Paired Kruskal-Wallis test adjusted for multiple comparisons was used for comparisons of the same treatment across age groups. Unpaired t-test and one-way ANOVA was used for statistical comparison where indicated. Only p values <0.05 are shown, and p<0.05 was considered statistically significant. Survival curves were created using the Kaplan and Meier method in Prism.

## Results

### Aging leads to hypergammaglobulinemia and appearance of autoantibodies in B10.S mice

To determine the relationship between age and autoantibody development in B10.S mice, the serum of mature (3 month old), adult (6 month old), middle aged (12 month old) and old (24 month old) male and female mice was tested for immunoglobulin levels, ANA, anti-chromatin, and anti-ENA5 autoantibodies. Mature mice did not show any evidence of ANA and only 1/19 adult mice was ANA positive; however, 34% (19/56) of middle aged mice were ANA positive with an average intensity score of 1.8 and 57% (27/47) of old mice developed ANA with an average score of 2.5 (Figure 1). ANA positivity occurred more frequently in females with 39% of middle aged female mice being ANA positive whereas 30% of males were ANA positive. Similarly, 65% of old female mice were ANA positive whereas only 50% of males were.

**Figure 1.**
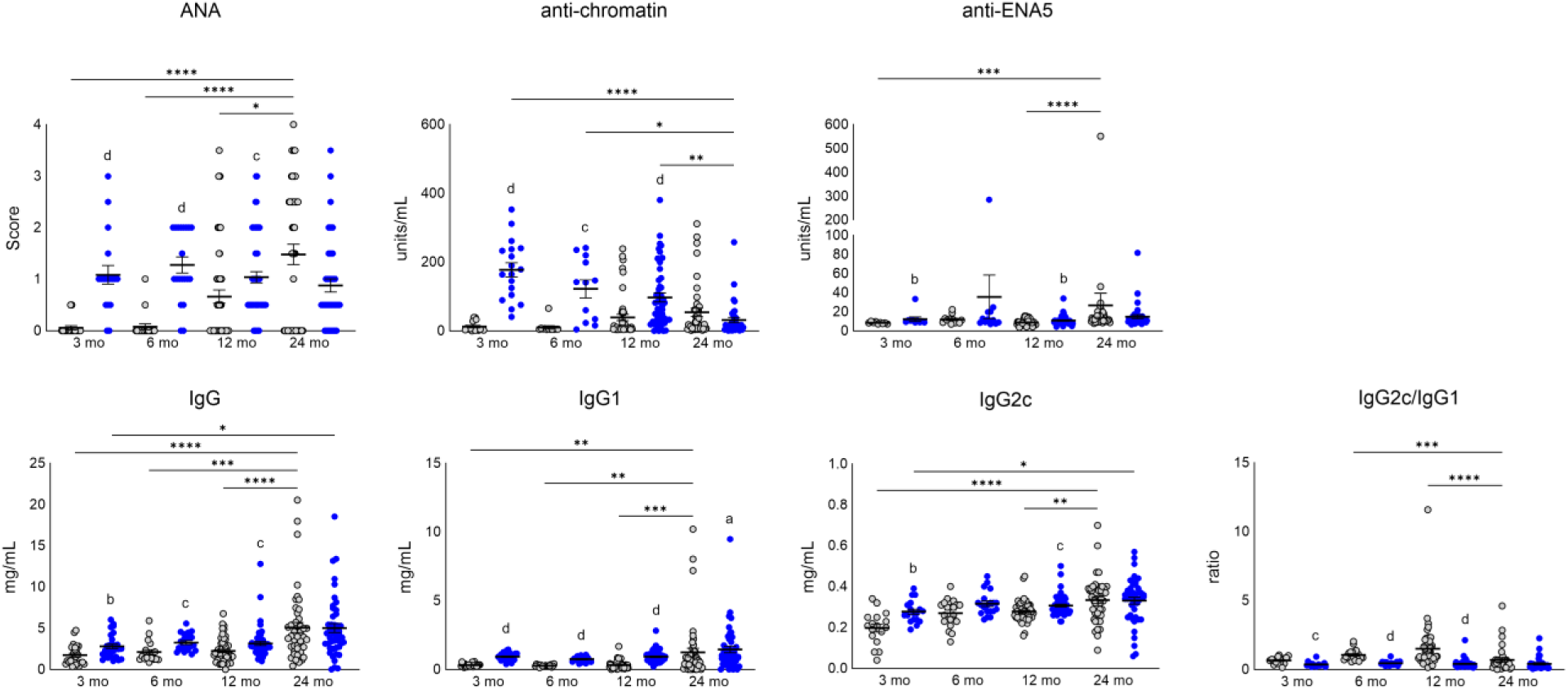
Age and HgCl_2_-induced changes in autoantibodies and hypergammaglobulinemia in B10.S mice. Sera assessment of B10.S mice at 3 (n=17/18), 6 (n=19/20), 12 (n=56/57) or 24 (n=48/44) months of age treated with PBS (gray) or HgCl_2_ (blue). Mice treatment and assays performed as described in Materials & Methods. Statistical comparisons between age groups were performed with Kruskal-Wallis tests. *p<0.05, **p<0.01, ***p<0.001, ****p<0.0001. Statistical comparisons of PBS and HgCl_2_ treatment groups were performed with Mann-Whitney U tests. a, p<0.05; b, p<0.01; c, p<0.001; d, p<0.0001.

Anti-chromatin antibodies, often found in systemic autoimmune diseases [51, 52], also increased with age (Figure 1). Anti-ENA5 antibodies, also found in patients with systemic autoimmune diseases, [53, 54], showed a slight increase with age but the majority of responses were below the mean+2 SD of mature mice (Figure 1). Total IgG, IgG1, and IgG2c increased with age (Figure 1). However, the ratio of IgG2c/IgG1 demonstrated an increase from mature to middle age; thereafter, the serum profiles of old age mice heavily favored IgG1 as their IgG2c/IgG1 ratio was significantly reduced in comparison to adult (p<0.001) and middle aged (p<0.0001) mice (Figure 1). These findings argue that aging is associated with enhanced humoral immunity including increased prevalence of autoantibodies.

### Presence of ANA is associated with hypergammaglobulinemia in middle and old age

Importantly, both middle aged and old mice consisted of ANA positive and negative animals, and this was associated with other serological differences. ANA positivity was associated with increased anti-chromatin antibodies in middle aged (p<0.0001) but not old mice (Figure 2). In ANA positive mice the ANA score was increased in old versus middle aged mice; however, this was not statistically significant. ANA had the strongest correlation with total IgG (r(137)=0.42, p<0.0001) (Supplemental Figure 2A). Increasing age was also correlated with increased anti-ENA5 responses (r(104)=0.47, p<0.0001). However, the lack of strong correlation between ANA and anti-ENA5 (r(104)=0.23, p<0.05) or anti-chromatin (r(104)=0.07, ns) autoantibody responses argues that neither response fully explains the increased intensity of the ANA response with age.

**Figure 2.**
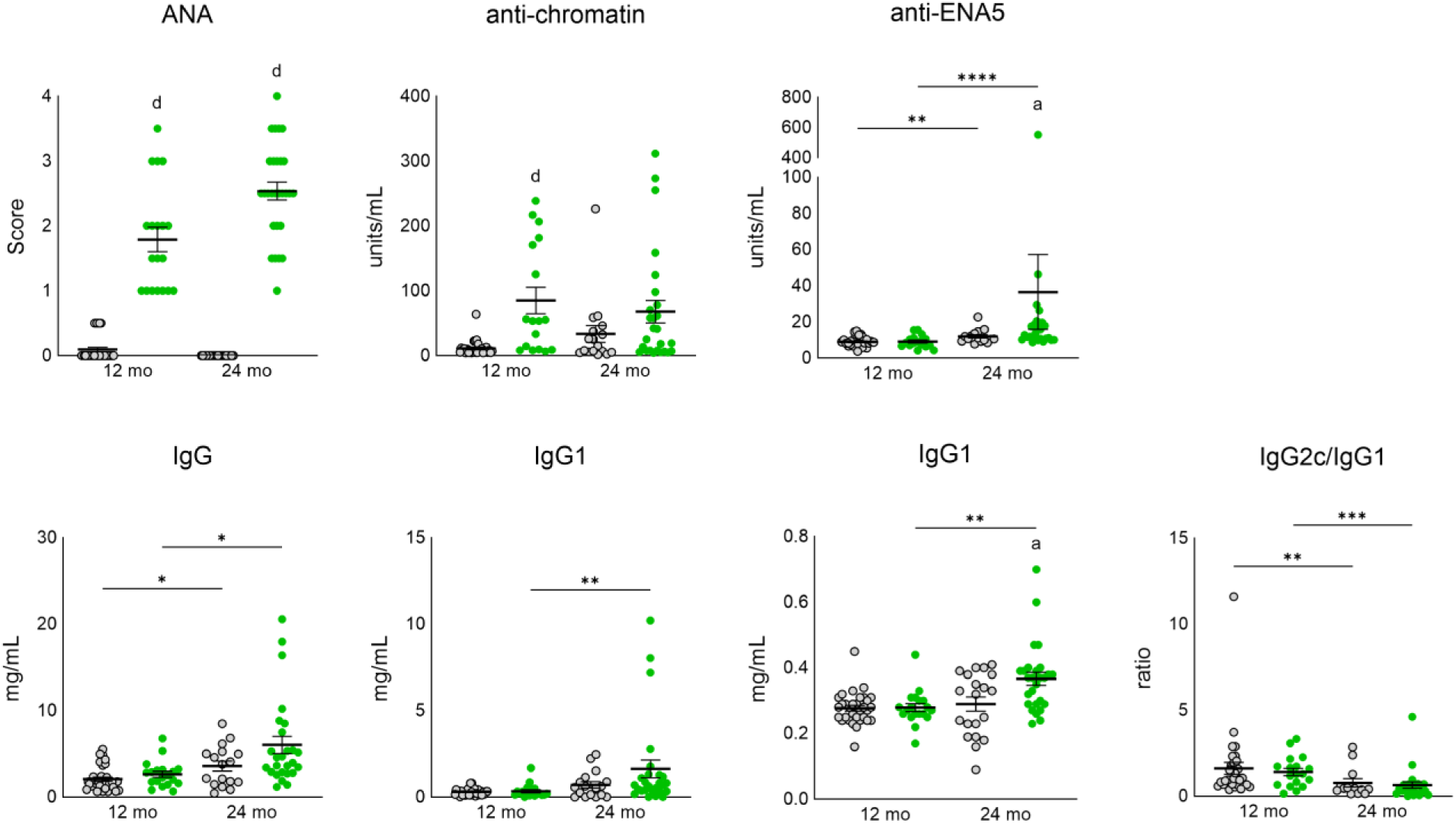
Comparison of humoral immune responses in ANA negative and positive middle aged and old age B10.S mice. Sera assessment of ANA negative **(gray)** and positive **(green)** 12 (n=32/19) and 24 (n=20/28) month old mice treated with PBS. Mice treatment and assays performed as described in Materials & Methods. Statistical comparisons between age groups were performed with Kruskal-Wallis tests. *p<0.05, **p<0.01, ***p<0.001, ****p<0.0001. Statistical comparisons of ANA negative and positive treatment groups were performed with Mann-Whitney U tests. a, p<0.05; b, p<0.01; c, p<0.001; d, p<0.0001.

These findings demonstrate that increasing age is associated with increasing presence of autoantibodies, especially ANA, and immunoglobulins. However, among middle aged and old mice two serum profiles develop. One group is positive for ANA and displays increased humoral immunity, while the other is negative for ANA and displays smaller elevations in immunoglobulins and other autoantibodies.

### HgCl_2_ exposure augments humoral immunity regardless of age

Mercury-induced autoimmunity (HgIA) is characterized by elevated levels of immunoglobulins and autoantibodies, especially an MHC-restricted anti-nucleolar autoantibody (ANoA) response [42]. However, the effect of age on these responses has not been examined. Mature mice showed increases in ANA (p<0.0001), anti-chromatin antibodies (p<0.0001), anti-ENA5 antibodies (p<0.01), total IgG (p<0.01), IgG1 (p<0.0001), and IgG2c (p<0.01) with exposure to HgCl_2_ compared to control mice exposed to PBS. Adult mice showed increases in ANA (p<0.0001), anti-chromatin antibodies (p<0.001), total IgG (p<0.01), and IgG1 (p<0.0001) with exposure to HgCl_2_, and middle aged mice showed significant increases in ANA (p<0.001), anti-chromatin antibodies (p<0.0001), anti-ENA5 antibodies (p<0.01), total IgG (p<0.001), IgG1 (p<0.0001), and IgG2c (p<0.001) with exposure to HgCl_2_ (Figure 1). Mercury exposure was associated with a significant reduction in the IgG2c/IgG1 ratio in mature (p>0.001), adult (p<0.0001), and middle aged (p<0.0001) mice, suggesting that while HgCl_2_ led to elevations in both IgG1 and IgG2c, IgG1 is the predominant response.

As expected, ANoA were induced by HgCl*_2_* in mature, adult and middle aged mice (p<0.0001), but although ANoA were induced in old mice (p<0.0001), the incidence was lower than for younger animals (Figure 3). Apart from ANoA, HgCl_2_ exposure was associated with elevated levels of IgG1 in old mice (p<0.05). This was a modest elevation compared to younger mice; however, the significance was increased by removing ANoA negative mice (p<0.001) (Supplemental Figure 3A). As was the case for younger mice, mercury exposure did not lead to a significant reduction in the IgG2c/IgG1 ratio (Figure 1), but removal of ANoA negative mice revealed a significant difference between PBS and ANoA positive old mice (Supplemental Figure 3B). Moreover, while HgCl_2_ exposure led to elevated anti-chromatin antibodies in mature, adult and middle aged mice, this induced response was not significantly elevated when compared to control mice in old age (Figure 1). Even after removing ANoA negative mice, HgIA was not associated with elevated anti-chromatin antibodies in old age mice. Interestingly, ANoA negative mice revealed significantly reduced levels of anti-chromatin antibodies when compared to control mice (p<0.05) (Supplemental Figure 3A), suggesting that failure to induce HgIA is associated with reductions in other autoantibodies.

**Figure 3.**
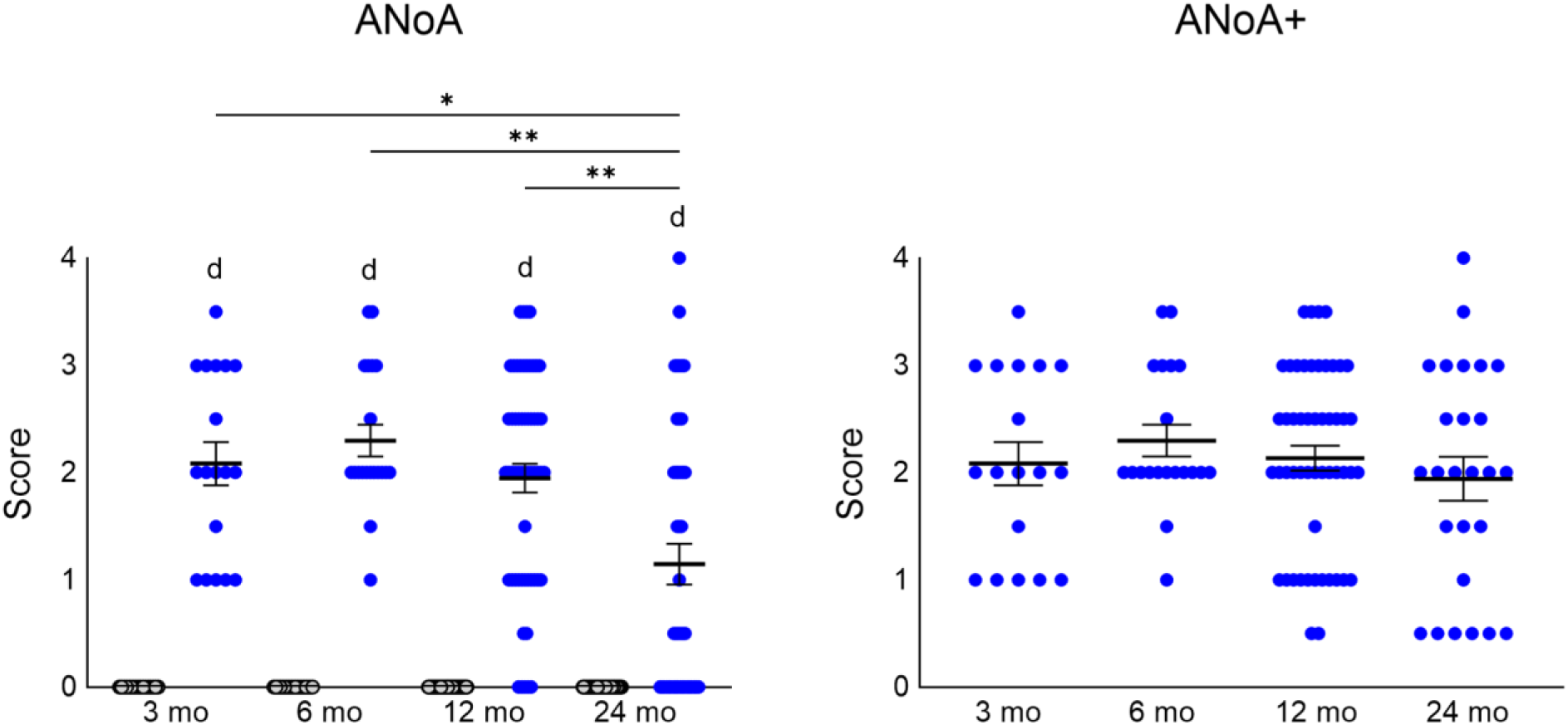
The ANoA response requires HgCl_2_-exposure but does not occur in all animals with age. Assessment of ANoA in B10.S mice at 3 (n=17/18), 6 (n=19/20), 12 (n=56/57) or 24 (n=48/44) months of age treated with PBS (gray) or HgCl_2_ (blue). Mice treatment and assays performed as described in Materials & Methods. Statistical comparisons between age groups were performed with Kruskal-Wallis tests. *P<0.05, **p<0.01, ***p<0.001, ****p<0.0001. Statistical comparisons of PBS and HgCl_2_ treatment groups were performed with Mann-Whitney U tests. a, p<0.05; b, p<0.01; c, p<0.001; d, p<0.0001.

### The incidence and severity of HgCl_2_-induced autoimmunity declines with age

Notably, only 59% (26/44) of old age mice were ANoA positive compared to mature, adult and middle aged animals where 100% (18/18), 100% (20/20), and 91% (52/57), respectively, of HgCl_2_ exposed mice were ANoA positive when sera was tested at a 1:100 dilution (Figure 3). However, comparison of ANoA positive mice revealed that age does not affect ANoA intensity (Figure 3). Upon retesting initially ANoA negative samples at a 1:40 dilution, 56% (10/18) of these samples had detectable ANoA, suggesting that while aging does not inhibit the production of mercury-induced ANoA, the strength of the humoral immune response does diminish with age. These findings argue that generation of ANoA in the genetically susceptible B10.S requires a specific stimulus other than physiological age. The data also shows that while HgCl_2_ exposure can induce robust ANoA responses at all ages, the intensity of this response is differentially affected by age.

Comparison of humoral immune responses in ANoA positive and negative HgCl_2_ exposed old age mice revealed increases in anti-ENA5 reactivity (p<0.05) (Figure 4). Moreover, removing mice with detectable ANoA when tested at a 1:40 dilution from the negative group resulted in ANoA positive (at 1:100 dilution) mice having significantly higher anti-chromatin antibodies (p<0.01) than ANoA negative mice (Supplemental Figure 3B). 38% (10/26) of ANoA positive mice were ANA positive and 39% (7/18) of ANoA negative mice were ANA positive (Figure 4), demonstrating that the spontaneous production of ANA is a poor indicator of mercury-induced ANoA. Significantly, ANoA positivity was linked to increases in total IgG1 (p<0.05), suggesting that the ability to produce HgCl_2_ specific ANoA in old age may reflect maintenance of T cell dependent B cell function.

**Figure 4.**
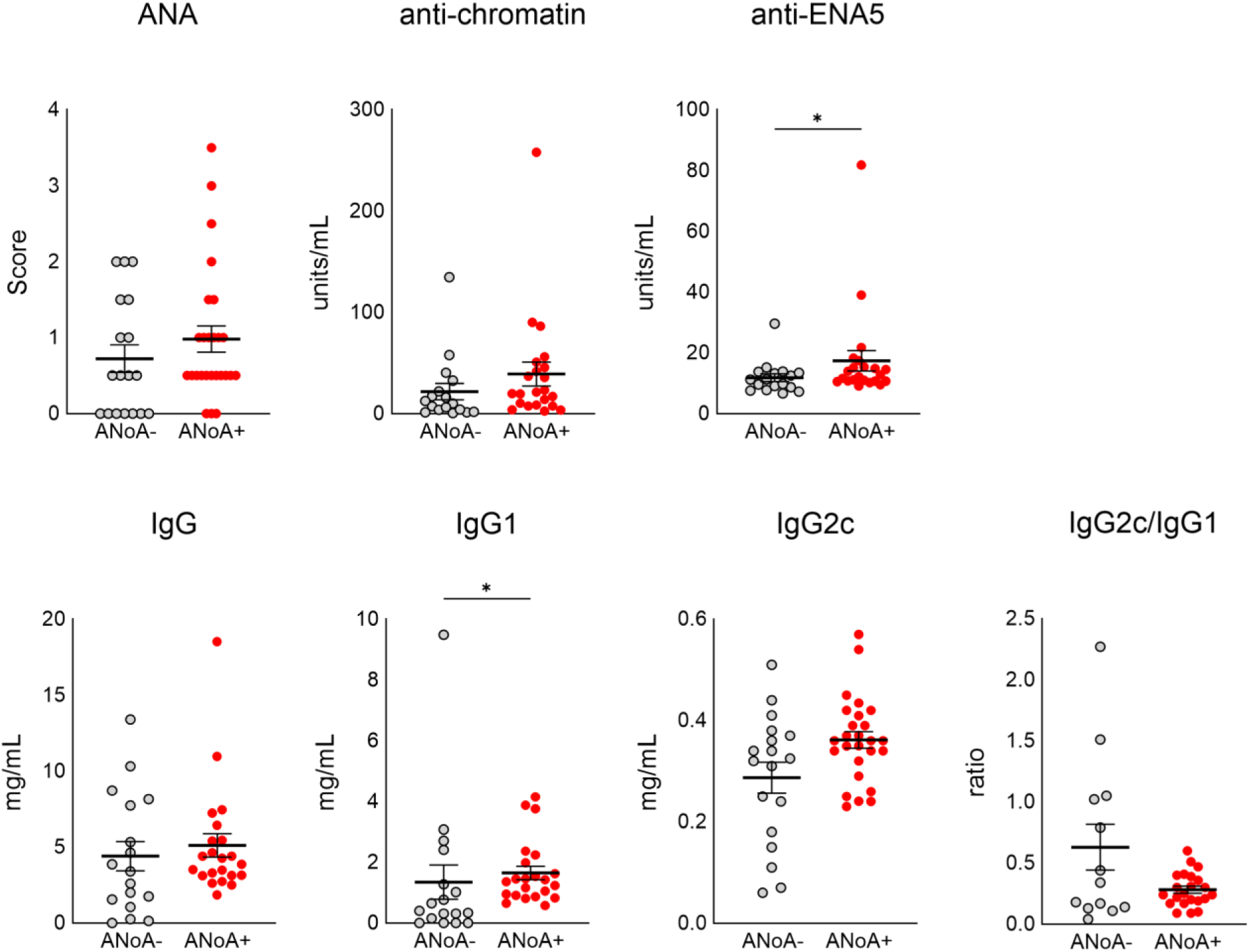
Comparison of humoral immune responses in ANoA negative and positive HgCl_2_ exposed old age mice. Sera assessment of ANoA negative (n=18) and ANoA positive (n=26) 24 month old mice treated with HgCl_2_. Mice treatment and assays performed as described in Materials & Methods. Statistical comparisons were performed with Mann-Whitney U tests. *P<0.05, **p<0.01, ***p<0.001, ****p<0.0001.

### Aging and HgCl_2_ effects on T cells

In mature age mice autoantibody development in HgIA is dependent upon CD4^+^ T cells [55] which proliferate in vitro [56, 57] and upregulate activation markers in vivo [58, 59] in response to HgCl_2_. Using multiparameter flow cytometry we evaluated if HgCl_2_ exposure resulted in age-related changes in splenic CD4^+^ (Figure 5) and CD8^+^ (Figure 5) T cell subsets.

**Figure 5.**
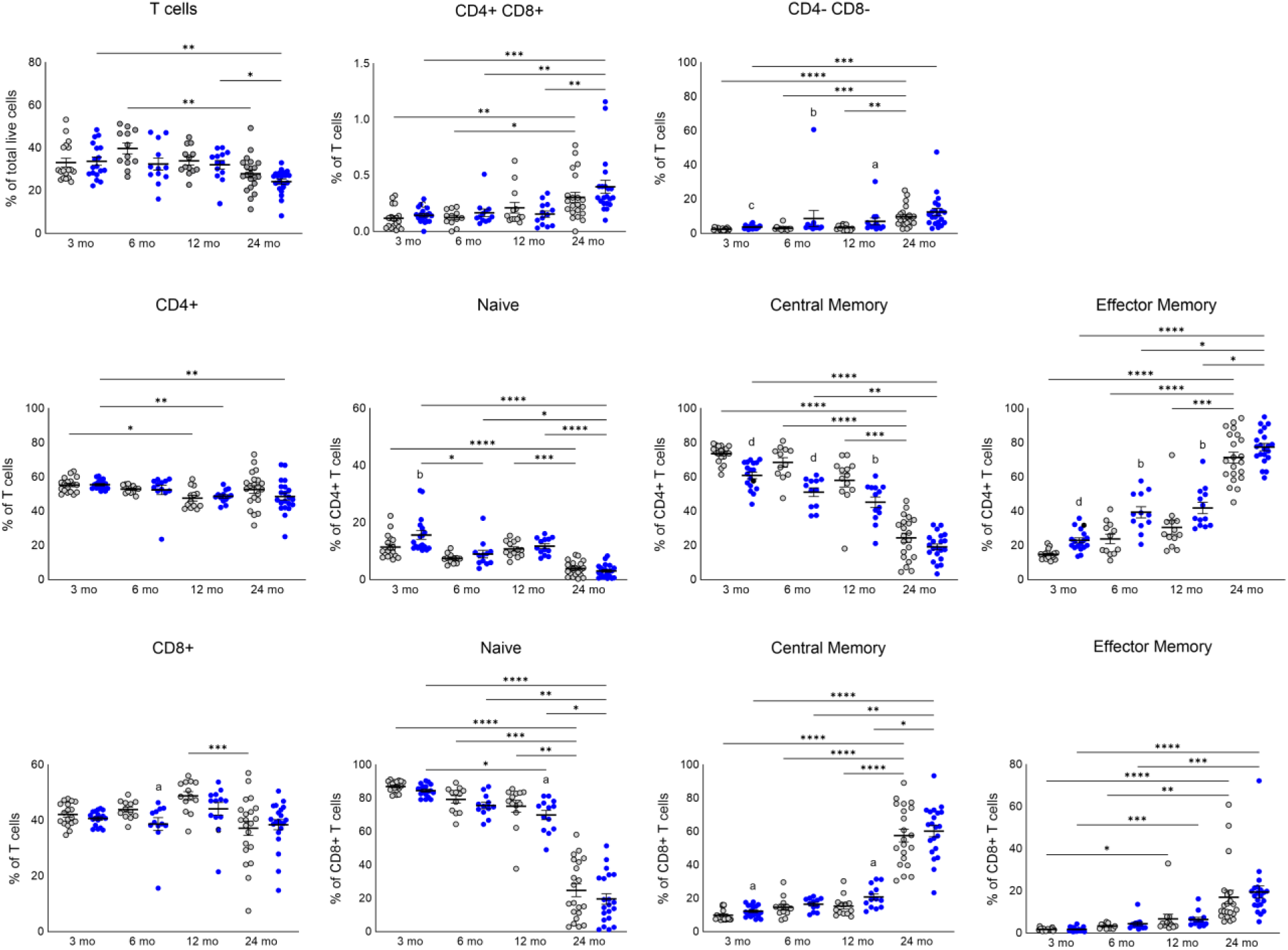
Changes in T cell subsets with age and HgIA. Multicolor flow cytometry on the spleens of B10.S mice at 3 (n=18/18), 6 (n=12/12), 12 (13/13) or 24 months (n=19/23) of age treated with PBS (gray) or HgCl_2_ (blue). Mice treatment and assays performed as described in Materials & Methods. Statistical comparisons between age groups were performed with Kruskal-Wallis tests. *P<0.05, **p<0.01, ***p<0.001, ****p<0.0001. Statistical comparisons of PBS and HgCl_2_ treatment groups were performed with Mann-Whitney U tests. a, p<0.05; b, p<0.01; c, p<0.001; d, p<0.0001.

The frequency of TCRβ^+^ cells declined with age (Figure 5 top panel), consistent with age-related decline of the T cell reservoir and adaptive immunity [60]. Unexpectedly, the frequencies of both double positive (CD4^+^CD8^+^) and double negative (CD4^−^CD8^−^) T cells increased with age as a percentage of total T cells. The expansion of double negative T cells was augmented by HgCl_2_ in mature (p<0.001), adult (p<0.01), and middle aged (p<0.05) mice.

### Aging and HgCl_2_ effects on CD4 T cells

The frequency of CD4^+^ T cells, as a percentage of TCRβ^+^ cells, was significantly reduced from mature to middle aged mice (p<0.05) (Figure 5 center panel). It does not appear that this reduction is maintained in old age mice treated with PBS; however, comparison of HgCl_2_-treated animals reveals a significant reduction of CD4^+^ T cells between mature and middle-aged mice (p<0.01) as well as mature and old age mice (p<0.01). Age-related decline of naïve and central memory (CM) CD4^+^ T cell subsets was observed, while the effector memory (EM) subset showed an increase with age. In old age mice, 71.4% (mean frequency) of CD4^+^ T cells had differentiated into EM cells. Many of these age-related changes were augmented by HgCl_2_ in mature, adult and middle aged mice. Mercury exposure in mature mice was associated with a reduction in the frequency of CM cells (p<0.0001) and increased frequencies of both naïve (p<0.01) and EM subsets (p<0.0001), suggesting that HgCl_2_ triggers the differentiation of CM cells into EM cells but has less influence on the conversion from naïve to CM. A reduction in CM and increase in EM subsets also occurred with mercury exposure in adult and middle aged mice; however, old age mice did not exhibit these HgCl_2_-induced changes and subset frequencies demonstrated high interindividual variability in both treatment and control groups. Thus, although CD4^+^ T cells differentiate with age, mercury exposure mimics these age-related changes when exposure occurs before old age.

### Aging and HgCl_2_ effects on CD8 T cells

The mean percentage of CD8^+^ T cells did not change linearly with age, despite there being a significant reduction from middle to old age (p<0.001) (Figure 5 bottom panel). Aside from a modest reduction in the frequency of CD8^+^ T cells in adult mice (p<0.05), HgCl_2_ did not influence CD8^+^ T cells as a percentage of total TCRβ^+^ cells. The percentage of naïve CD8^+^ T cells were decreased in old age but, in contrast to CD4^+^ T cells, CM CD8^+^ T cells increased with age (Figure 5 bottom panel). The percentage of CD8^+^ EM T cells also increased in old age. These age-related changes in CD8^+^ T cell subsets were not significantly affected by exposure to HgCl_2_. Comparison of HgCl_2_-induced changes within each age group showed that HgCl_2_ exposure modestly decreased the percentage of naïve CD8^+^ T cells in middle aged mice and increased CM CD8^+^ T cells in mature and middle aged mice (all p<0.05). As with CD4^+^ T cells, aging was associated with CD8^+^ T cell differentiation and changes in CD8^+^ T cell subsets were not affected by mercury exposure. Consistent with HgIA being a T helper-dependent response, HgCl_2_ exposure led to more subtle changes of CD8^+^ T cell subsets within each age group.

Together, these observations confirm the increased variability of T cell subset percentages in old age, supporting the global changes that occur in T cell populations with age [60, 61]. While mercury exposure was not associated with any significant changes to the frequencies of T cell subsets in old age mice, the interindividual variability observed for both treatment and control groups could obscure mercury-induced changes. The data provides the novel observation that mercury exposure can induce changes in T cell populations in mature, adult and middle aged mice suggesting that mercury may modulate T cell function irrespective of age.

### Germinal center CD4 T cells are increased with HgCl_2_ exposure

Germinal center (GC) CD4^+^ T cells are a distinct subset of differentiated T follicular helper (T_FH_) cells that reside in germinal centers and have an enhanced B cell help capacity [62]. Zheng et al. have evidenced that this is a population of antigen-specific CD4^+^ T cells that migrate to the germinal center rather than a distinct lineage [63]. GC T_FH_ cells have been identified based on their coexpression of CXCR5 which allows B cell follicle homing, and GL7 which is typically used to identify GC B cells. We determined this population as a subset of EM CD4^+^ T cells that were GL7^+^CD38^+^. As a percentage of total CD4^+^ T cells, GL7^+^CD38^+^ cells increased linearly with age. However, as a percentage of EM CD4^+^ T cells, the GC subset followed a nonlinear trajectory that suggested a relative decline after middle age (Figure 6 bottom panel). However, this is occurring within the context of age-related differentiation suggesting that the differentiation of CD4^+^ T cells with age does not lead to proportional increases of T helper cells in the germinal center. Mercury exposure led to profound increases in this population as a percentage of EM cells and total CD4^+^ T cells among mature (p<0.0001), adult (p<0.0001), and middle aged (p<0.001) mice; however, this elevation was not observed in old age mice (Figure 6 bottom panel). Even when ANoA negative mice were removed, this population was not significantly elevated in old mice with HgIA.

**Figure 6.**
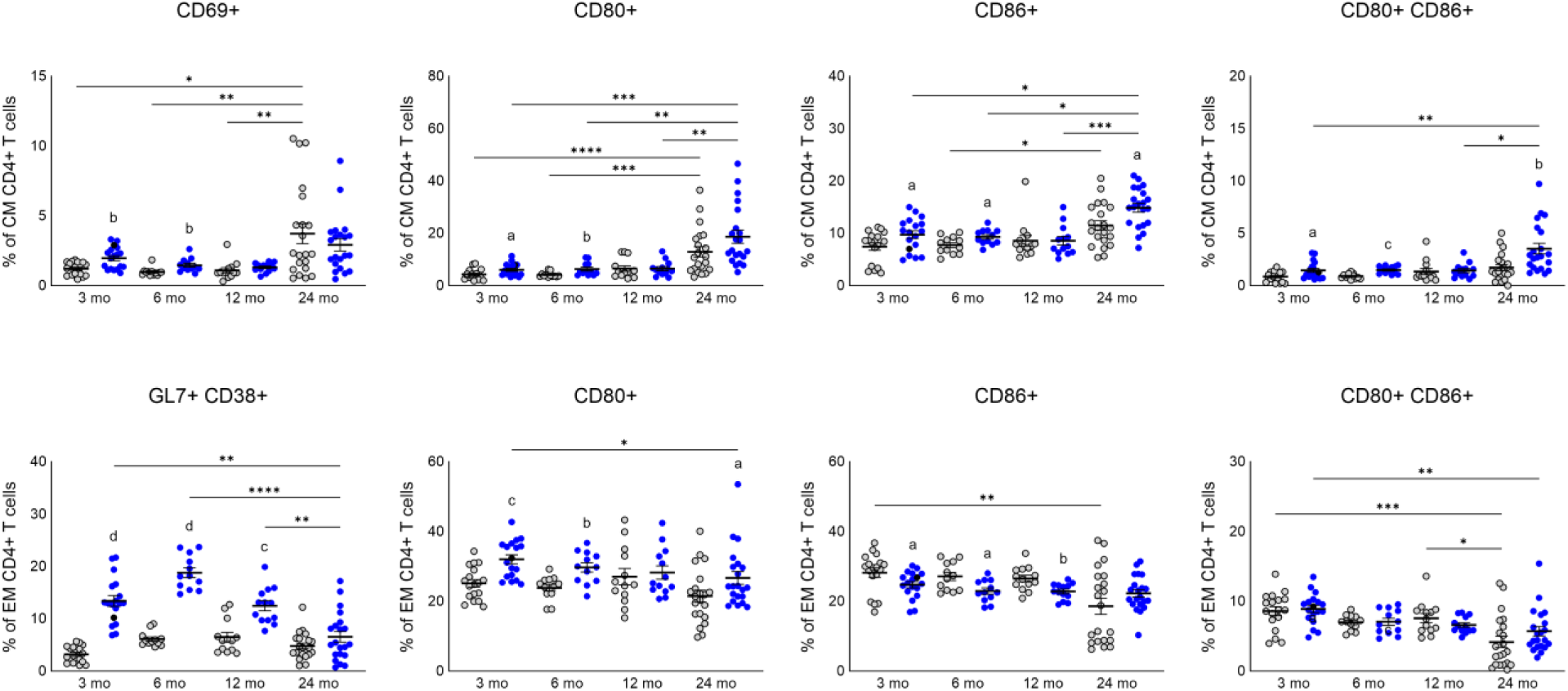
Activation of CD4^+^ T cell subsets with age and HgIA. Multicolor flow cytometry on the spleens of B10.S mice at 3 (n=18/18), 6 (n=12/12), 12 (13/13) or 24 (n=19/23) months of age treated with PBS (gray) or HgCl_2_ (blue). Mice treatment and assays performed as described in Materials & Methods. Statistical comparisons between age groups were performed with Kruskal-Wallis tests. *P<0.05, **p<0.01, ***p<0.001, ****p<0.0001. Statistical comparisons of PBS and HgCl_2_ treatment groups were performed with Mann-Whitney U tests. a, p<0.05; b, p<0.01; c, p<0.001; d, p<0.0001.

After identifying this population as being associated with HgIA, we conducted a follow-up study in middle aged mice to confirm CXCR5 expression. Our results revealed that the vast majority of GL7^+^ CD4^+^ T cells were CXCR5^+^, confirming a T_FH_ phenotype. Although we did not find a significant difference in GL7^+^CXCR5^+^ cells as a percentage of EM CD4^+^ T cells (p=0.08), there was a significant difference in the total number of GC T_FH_ (p<0.01) (Figure 7). There was a relative reduction in the frequency of GL7^−^CXCR5^+^ cells and no change in the frequency of CXCR5 expression among EM CD4^+^ T cells. Furthermore, there was not a significant elevation of CM T_FH_. This suggests that mercury is influencing the migration of EM T_FH_ into the germinal center rather than inducing the selective expansion of an independent lineage.

**Figure 7.**
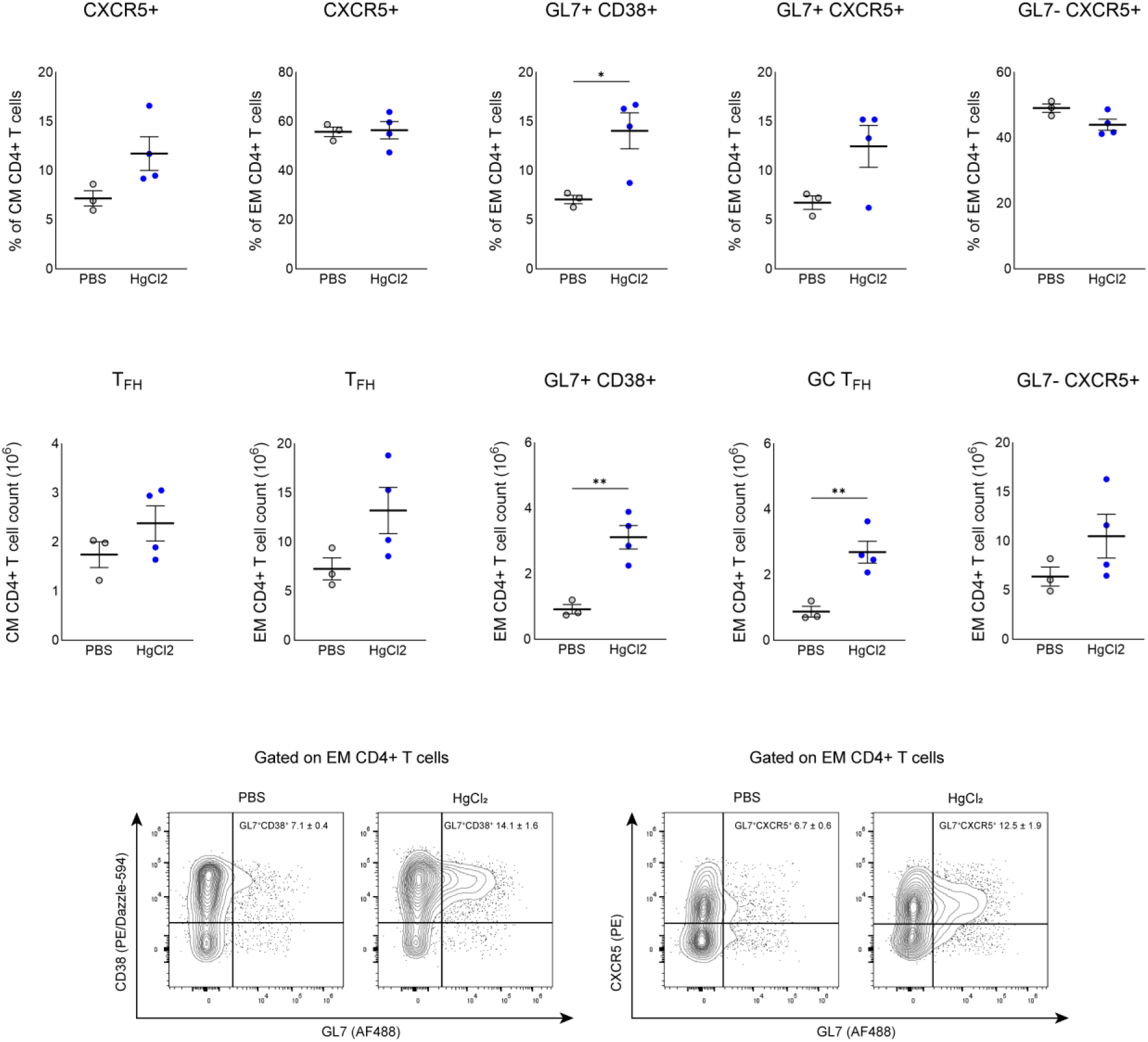
Germinal center T follicular helper cells with HgIA. Multicolor flow cytometry on the spleens of B10.S mice at 12 months of age treated with PBS (n=3) (gray) or HgCl_2_ (n=4) (blue). Mice treatment and assays performed as described in Materials & Methods. Statistical comparisons between PBS and HgCl_2_ treatment groups were performed with unpaired t-tests (black *). *P<0.05, **p<0.01, ***p<0.001, ****p<0.0001.

### B cell subset profiles change with age

Multiparameter flow cytometry of splenic B cells identified significant age-related changes in B cell subsets reflected primarily by the increased variation of subset frequencies in old age mice (Figure 8). The population of CD19^+^ B cells in mature mice was comprised of 8.0% transitional, 2.9% B1 and 87.4% B2 (mean frequencies) cells. These frequencies are relatively maintained for adult and middle aged mice; however, transitional B cells are depleted, B2 cells are reduced and B1 cells are significantly increased in old age mice. Notably, B1 B cells went from representing a small minority of B cells in mature, adult and middle aged mice to 37.3% (mean frequency) of the total CD19^+^ B cell population in old mice. Considering the standard deviation of B1 B cells was 29.6% in old mice, B1 cells represented the majority of B cells in a considerable number of old age mice. This finding is consistent with the impairment of B2 cell development, the expansion of B1 cells, and a shift from a predominantly adaptive immune response to innate immunity that occurs with age [64, 65].

**Figure 8.**
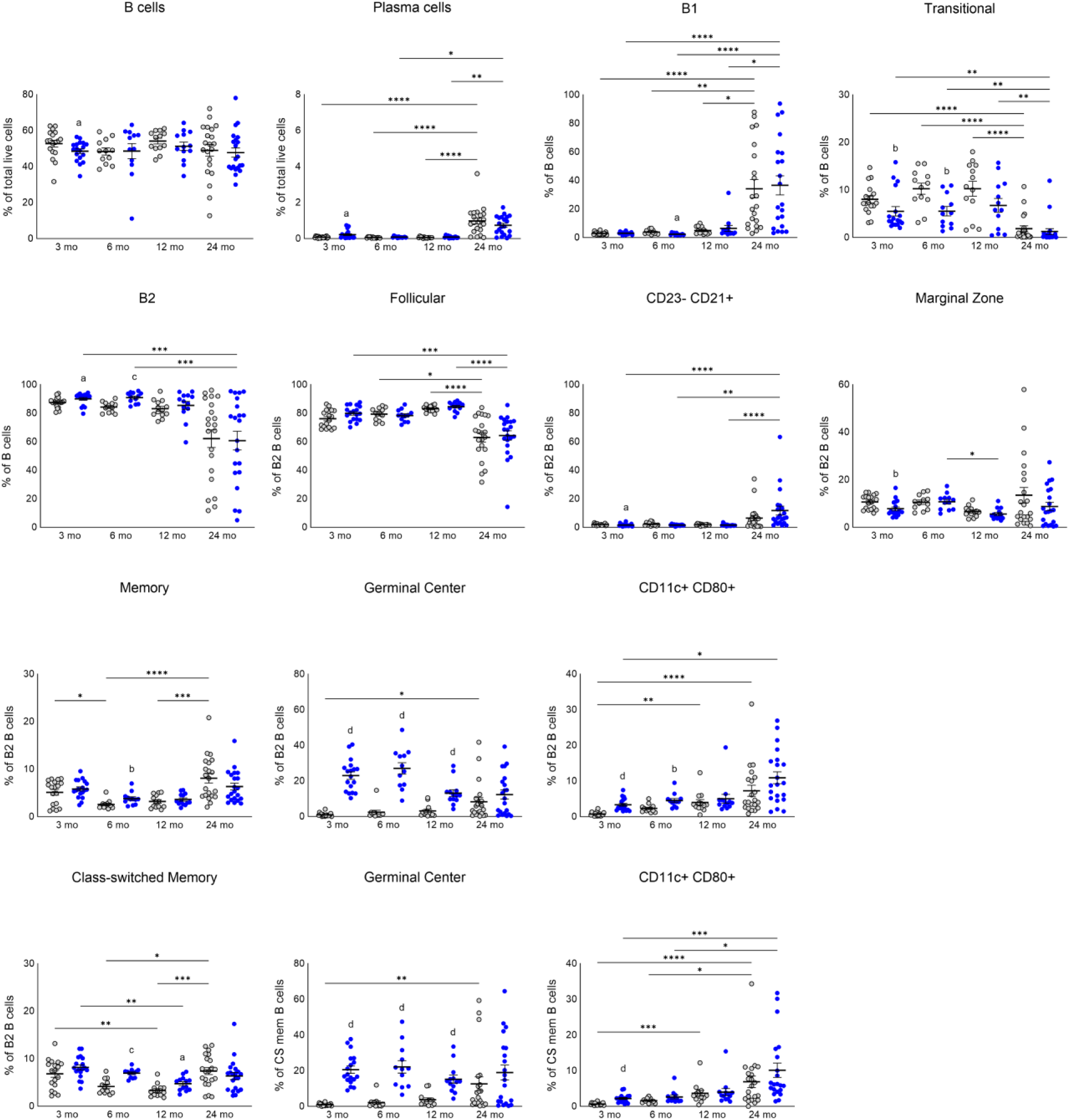
B cell subset changes with age and HgIA. Multicolor flow cytometry on the spleens of B10.S mice at 3 (n=18/18), 6 (n=12/12), 12 (13/13) or 24 (n=19/23) months of age treated with PBS (gray) or HgCl_2_ (blue). Mice treatment and assays performed as described in Materials & Methods. Statistical comparisons between age groups were performed with Kruskal-Wallis tests. *P<0.05, **p<0.01, ***p<0.001, ****p<0.0001. Statistical comparisons of PBS and HgCl_2_ treatment groups were performed with Mann-Whitney U tests. a, p<0.05; b, p<0.01; c, p<0.001; d, p<0.0001.

While the frequencies of follicular (FO), marginal zone (MZ) and memory B cells changed across age groups, age-related changes were not linear. Surprising changes in B2 B cell subtypes were observed in old age mice, primarily the emergence of an intermediate population that appeared between the MZ and memory populations on a CD23 versus CD21 plot, expressing moderate levels of CD21 but lacking CD23 expression. Germinal center (GC) B cells and plasma cells were increased in old age mice, likely resulting from lifetime exposure to foreign- and self-antigens.

### B cell subset profiles change with HgCl_2_ exposure

Although mercury exposure is linked to autoantibodies [6, 42], very little is known about the B cell subsets producing the autoantibody response. Apart from a modest increase in the frequency of the intermediate B2 population described above, B cell populations were little changed by HgCl_2_ exposure in old mice (Figure 8). Mercury exposure was associated with a significant increase in the frequency of B2 B cells in mature (p<0.05) and adult (p<0.001) mice. This was reflected by a reduction in the frequency of transitional B cells in mature (p<0.01) and adult (p<0.01) mice and a reduction in the frequency of B1 cells in adult mice (p<0.05). Although there was not a significant difference between the frequencies of transitional B cells in middle aged mice, interindividual variability increased in middle age and a few control mice demonstrated a depleted population. In old age mice, mercury was not able to further reduce the frequency of transitional B cells beyond the already depleted population observed in control mice. Mercury-related changes in B2 B cell subtypes were limited and inconsistent across age groups, mercury exposure was associated with a reduction in MZ B cells in mature mice and an increase in memory B cells in adult mice (both p<0.01).

The most profound effect of mercury exposure was demonstrated by the increased frequency of GC B cells in mature, adult and middle aged mice (all p<0.0001). As for GC T cells, mercury did not lead to significant elevations of GC B cells above controls in old age mice, consistent with the reduction of germinal center responses with age [66]. These findings, together with the preponderance of B2 B cells and increased EM CD4^+^ T cells, argue that mercury exposure induces a more effective T-B cell response in younger mice. Furthermore, mercury induced profound increases in germinal center B and T cells, changes that were not observed in DBA/2 mice that do not produce ANoA upon mercury exposure (data not shown), demonstrating that germinal center formation is central to the development of HgIA.

### Age associated B cells (ABCs) are increased with HgCl_2_ exposure

ABCs accumulate with age but arise earlier and more prominently in autoimmune-prone mouse strains [30], and are associated with autoantibodies including anti-chromatin autoantibodies [29, 30]. A phenotypically similar population in humans, known as atypical B cells, have been associated with immune responses to vaccination, infection, and autoimmune disease [67]. Studies of ABCs have used a variety of gene expression markers in different contexts [30, 68]. More recent studies using single-cell RNA sequencing have determined ABCs are a highly conserved memory B cell lineage with shared transcriptomes across different contexts [67]. Analysis of the scRNA-seq data to date has identified CD11c and T-bet are among the most consistently upregulated genes for human and murine ABCs [67]. We determined ABCs as a fraction of memory (excluding follicular, transitional, and marginal zone) CD19^+^B220^+^ (B2) B cells that were CD11c^+^CD80^+^CD21^−^CD23^−^ [29, 68]. These cells increased with age, as a percentage of memory B2 B cells, rising from 0.73% in mature to 6.3% in old age mice (mean frequencies) (p<0.0001). Mercury-induced increases were observed in mature (p<0.0001) and adult (p<0.01) mice (Figure 7). Mercury exposure did not lead to a statistically significant expansion of ABCs in middle aged or old mice beyond the levels observed in control mice.

While CD11c is broadly considered to be a reliable marker for the identification of ABCs, a significant population of CD11c^−^T-bet^+^ memory B cells have been identified in both mice and humans [69]. Because T-bet is widely regarded as the hallmark transcription factor of ABCs, we conducted a follow-up study to examine T-bet expression in splenic memory B cells following mercury exposure of mature, adult and middle aged mice. Consistent with the literature, all CD11c^+^ memory B cells highly expressed T-bet in addition to a population of CD11c^−^T-bet^+^ cells. Significantly, this population greatly outnumbered CD11c^+^T-bet^+^ memory B cells and expanded to a much greater extent in response to mercury. Consistent with our original study, T-bet^+^ memory B cells with and without CD11c expression were elevated in middle aged control mice (Figure 9). Furthermore, mercury exposure induced the most profound expansion of ABCs in adult mice, reflected by significantly greater percentages and total cell numbers when compared to mature and middle aged mice exposed to mercury. Mercury did not induce a significant elevation of CD11c or T-bet expression as a percentage of memory B cells in middle aged mice; however, these subsets did significantly increase in number. These findings demonstrate the expansion of ABCs in younger mice in the context of xenobiotic-induced autoimmunity.

**Figure 9.**
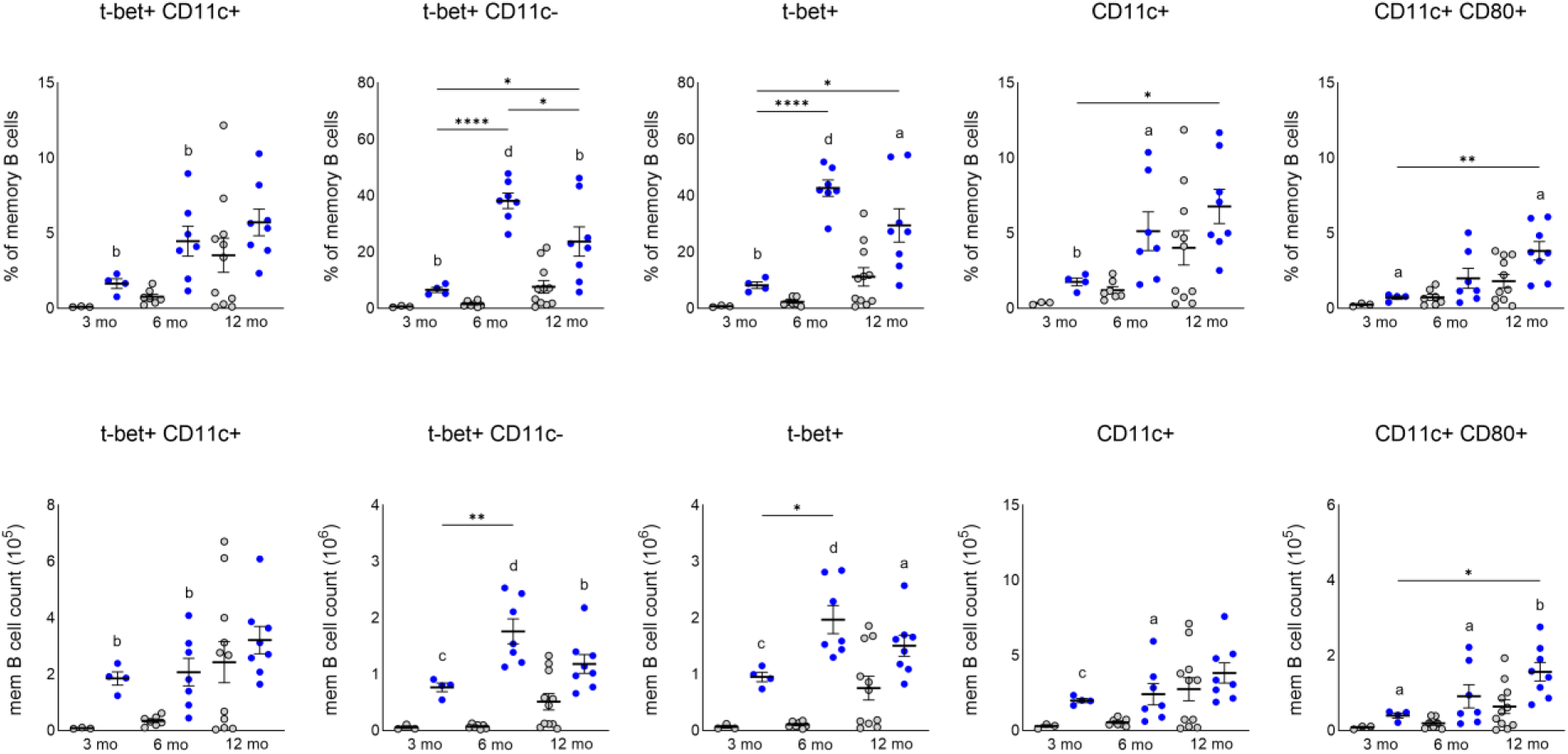
Age-associated B cells with age and HgIA. Multicolor flow cytometry on the spleens of B10.S mice at 3 (n=3/4), 6 (n=7/7) or 12 (11/8) months of age treated with PBS (gray) or HgCl_2_ (blue). Mice treatment and assays performed as described in Materials & Methods. Statistical comparisons between age groups were performed with one-way ANOVA. *P<0.05, **p<0.01, ***p<0.001, ****p<0.0001. Statistical comparisons between PBS and HgCl_2_ treatment groups were performed with unpaired t-tests. a, p<0.05; b, p<0.01; c, p<0.001; d, p<0.0001.

### Non T or B cell subset profiles change with age

Assessment of splenic cells in B10.S mice at progressive phases of life revealed the expansion of non T or B cells with age. As a percentage of total splenocytes, non T or B cells increased moderately from adult to old age (p<0.05) (Figure 10). Cell type analysis indicates that this expansion is driven primarily by the elevation of granulocytes in old mice. This observation is consistent with an aging phenotype driven by the production of granulocytes [70] that leads to the chronic innate immune activation characteristic of inflammaging [21, 71]. The elevation of granulocytes as a percentage of myeloid cells was reflected by corresponding decreases in the frequencies of monocytes (Figure 10). We identified a population of monocytes that expressed activation and antigen presentation markers CD38^+^CD80^+^CD86^+^, the frequency of which was not affected by age in PBS-treated mice. The frequency of dendritic cells demonstrated few age-related changes apart from the increased interindividual variability observed in old mice.

**Figure 10.**
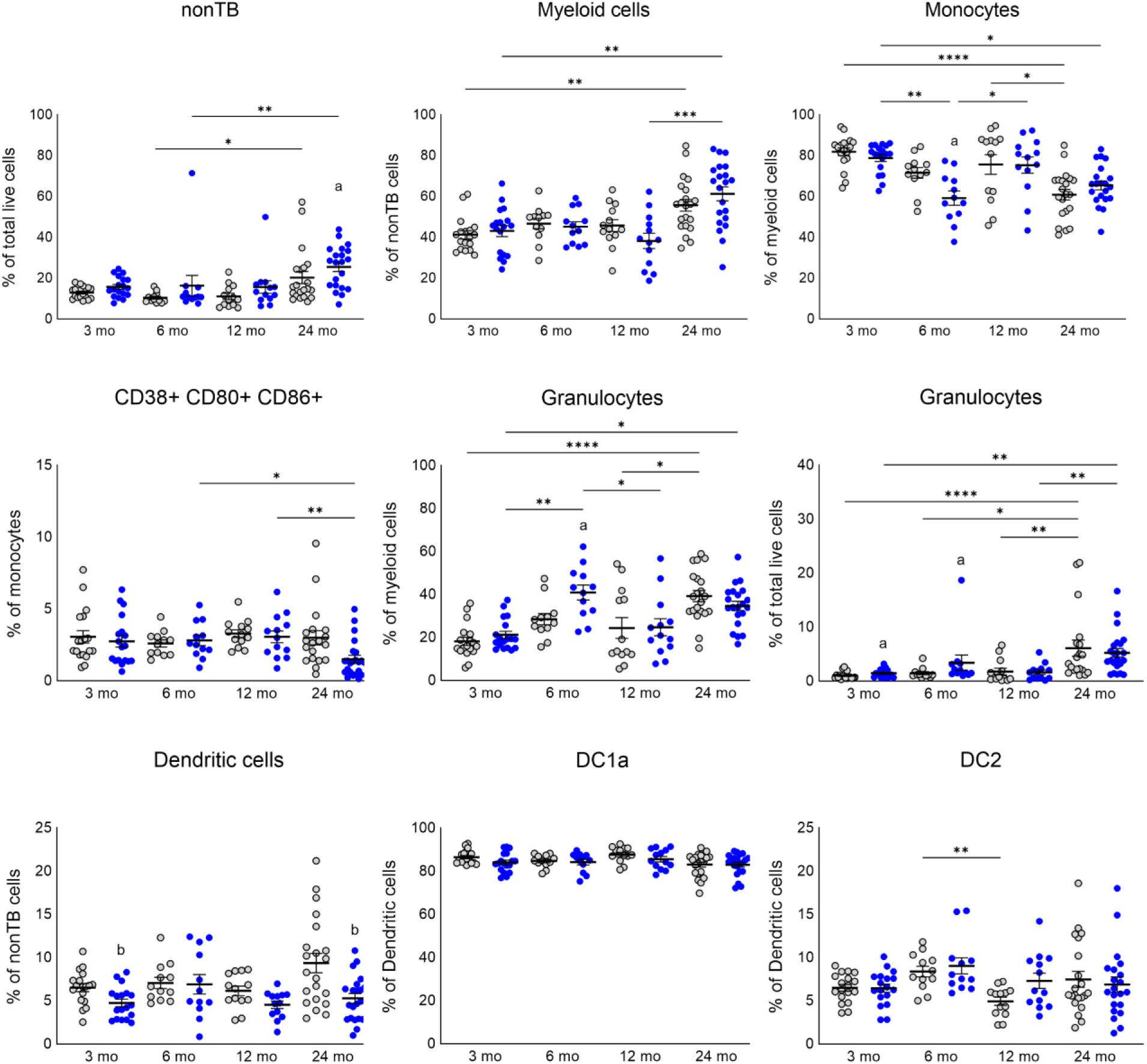
non T or B cell type changes with age and mHgIA. Multicolor flow cytometry on the spleens of B10.S mice at 3 (n=18/18), 6 (n=12/12), 12 (13/13) or 24 (n=19/23) months of age treated with PBS (gray) or HgCl_2_ (blue). Mice treatment and assays performed as described in Materials & Methods. Statistical comparisons between age groups were performed with Kruskal-Wallis tests. *P<0.05, **p<0.01, ***p<0.001, ****p<0.0001. Statistical comparisons of PBS and HgCl_2_ treatment groups were performed with Mann-Whitney U tests. a, p<0.05; b, p<0.01; c, p<0.001; d, p<0.0001.

### HgCl_2_ exposure is associated with increased innate immune cells

In comparison to T and B cell subsets, HgCl_2_ exposure was associated with few changes to non T or B cells. In mature and adult mice, mercury exposed mice had significantly elevated granulocytes as a percentage of total live cells (both p<0.05). This elevation was reflected by reduced populations of monocytes and dendritic cells. In old age mice, mercury augmented a significant elevation of non T or B cells (p<0.05) (Figure 10). However, the frequencies of CD38^+^CD80^+^CD86^+^ monocytes and dendritic cells were significantly reduced (both p<0.01), suggesting that the expansion of non T or B cells is not driven by an increase in antigen presenting cells (Figure 10). In sum, mercury induces the elevation of granulocytes and other innate immune cells which has the potential to produce inflammation that resembles early aging.

### Presence of ANA in middle-aged mice is associated with changes in T and B cell subsets

The presence of ANA in PBS-treated middle aged and old mice provides an opportunity to identify associated changes in T and B cell subsets. Unexpectedly, few differences were found in old age mice. However, the degree of interindividual variability could be obscuring changes in cell frequencies that are associated with the spontaneous production of ANA in old age.

Comparison of spleen cell profiles between ANA negative and positive middle aged mice revealed several changes that are associated with the spontaneous production of ANA. Significantly, many of the observed changes resemble those that occur with exposure to HgCl_2_. Double negative T cells were elevated among ANA positive mice but, unlike with mercury exposure, double positive T cells were also elevated (both p<0.05). Central memory CD4^+^ T cells were significantly reduced among ANA positive middle aged mice (p<0.05) (Figure 11); however, this was not reflected by a significant elevation of effector memory cells. Furthermore, T helper subsets demonstrated activation markers that mirrored observations made of mercury exposed animals; specifically, a significant elevation of GL7^+^CD38^+^ effector memory cells (p<0.05). Moreover, the importance of germinal centers for the production of autoantibodies is further evidenced by the elevation of memory (p<0.01) and germinal center (p<0.05) B cell frequencies among ANA positive mice. Additionally, ANA positive mice demonstrated reduced frequencies of transitional B cells (p<0.05) and increased B2 B cells (p<0.01). Spontaneous ANA production was also associated with changes in non T or B cell subsets, namely, reduced frequencies of dendritic cells and monocytes (both p<0.05) (Figure 11 bottom panel); however, these reductions were not reflected by an increase in granulocytes.

**Figure 11.**
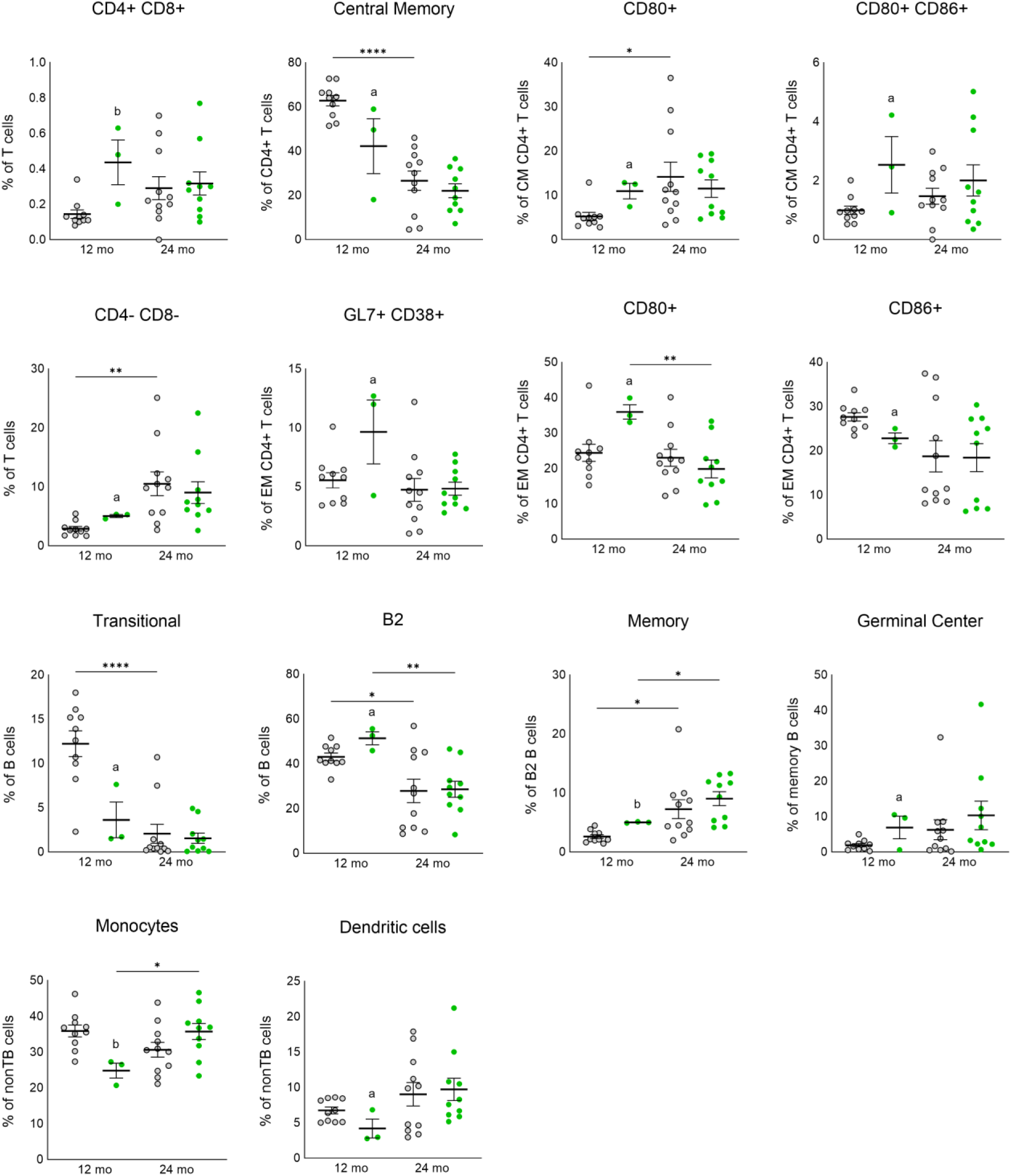
Spontaneous ANA is reflected in spleen cell populations in middle aged mice. Multicolor flow cytometry on the spleens of ANA-(gray) and ANA+ (green) 12 (10/3) and 24 (11/10) month old B10.S mice treated with PBS. Mice treatment and assays performed as described in Materials & Methods. Statistical comparisons between age groups were performed with ANOVA tests. *P<0.05, **p<0.01, ***p<0.001, ****p<0.0001. Statistical comparisons of PBS and HgCl_2_ treatment groups were performed with unpaired t-tests. a, p<0.05; b, p<0.01; c, p<0.001; d, p<0.0001.

In sum, the data reveals that similar cellular changes underly induced and spontaneous autoantibody production. This suggests that a common mechanism drives the development of spontaneous and induced autoimmunity.

### Presence of HgCl_2_-induced ANoA in old mice is associated with changes in immune cell subsets

The restriction of ANoA to HgCl_2_ exposure allowed examination of immune cell subsets linked to successful induction of this autoantibody response in old age. Comparison of the percentage of T, B and myeloid cell subsets between ANoA positive and negative old mice identified few significant differences (Figure 12). However, group numbers in relation to the high degree of deviation may be obscuring the detection of significant differences. Consistent with germinal center formation being central to HgIA, ANoA negative mice had significantly reduced germinal center B cells as a percentage of total B cells (p<0.05). Furthermore, ANoA negative mice demonstrated elevated percentages of non T or B cells (p<0.05), which may indicate hyperactive innate immunity characteristic of an aged immune system [21, 71]. Few significant differences were found for cell frequencies between PBS and HgCl_2_ treated old mice, and even when ANoA negative mice were removed, few significant differences were found between the PBS and ANoA positive groups. ANoA negative mice had elevated spleen weights (p<0.05). Therefore, in addition to the differences found for percentages, there may be differences in the number of cells within different cell types. Ultimately, while further research is needed to make definitive conclusions, the data suggests that the shift from adaptive to innate immunity that occurs in old age reduces the capacity to form germinal centers and produce autoantibodies in response to mercury exposure.

**Figure 12.**
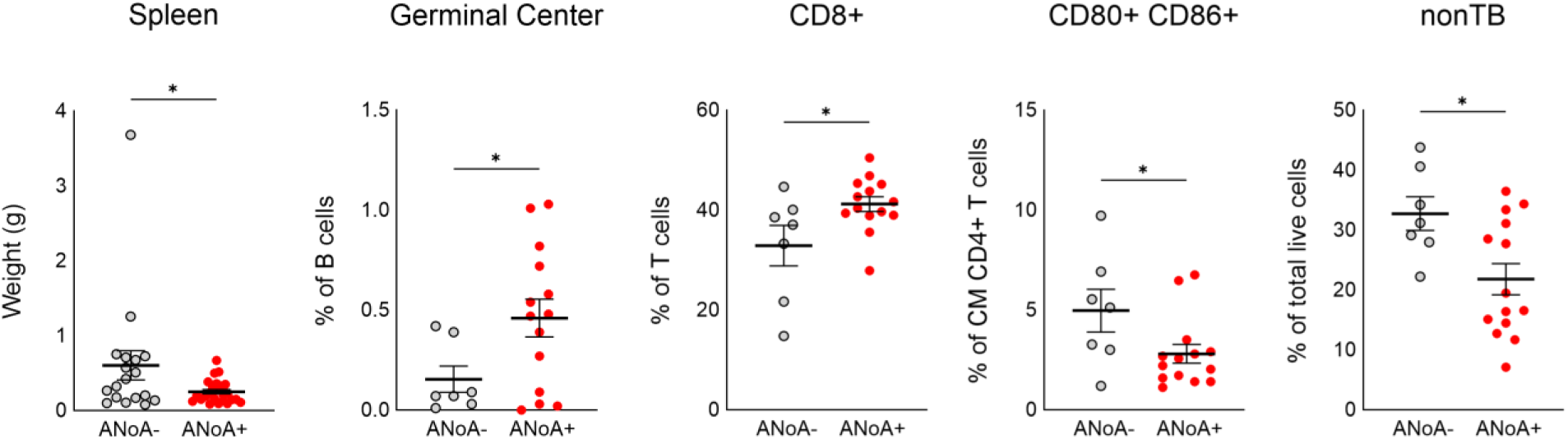
ANoA- and ANoA+ responses to HgCl_2_ are reflected in spleen cell populations in old mice. Multicolor flow cytometry on the spleens of ANoA-(n=7) and ANoA+ (n=16) 24 month old B10.S mice. Mice treatment and assays performed as described in Materials & Methods. Statistical comparisons between ANoA negative and positive mice were performed with unpaired t-tests. *P<0.05, **p<0.01, ***p<0.001, ****p<0.0001.

## Discussion

The findings of this study provide critical insights into the complex interplay between aging and xenobiotic-induced autoimmunity, specifically mercury-induced autoimmunity (HgIA) in B10.S mice. Our results demonstrate that while aging is associated with an increased prevalence of spontaneous autoantibodies, such as anti-nuclear antibodies (ANA), the capacity for xenobiotic-induced autoantibody responses, particularly anti-nucleolar autoantibodies (ANoA), diminishes in old age. This observation is significant, as it highlights a divergence between spontaneous and xenobiotic-induced autoimmune responses in the context of aging, offering novel perspectives on age-related immune dysregulation and its interaction with environmental exposures.

Our study utilized a robust sample size, substantially larger than that typically employed in mouse studies, which enabled the detection of differential immune responses in old age mice within an inbred strain. Among this cohort, only 59% of old mice developed a significant ANoA titer upon mercury exposure, compared to near-universal positivity in younger cohorts. This differential response among genetically identical mice underscores the impact of aging on immune variability, a phenomenon that would likely be exponentially amplified in genetically diverse populations or mouse models. The data confirm that aging in B10.S mice is associated with hypergammaglobulinemia and a progressive increase in ANA prevalence, consistent with prior reports of age-related subclinical autoimmunity in humans and mice [20, 23, 27, 37, 39]. Specifically, 34% of middle-aged and 57% of old age mice exhibited ANA positivity, with a pronounced female bias, aligning with known sexual dimorphism in autoimmune susceptibility [34, 35]. The rise in anti-chromatin antibodies and total IgG levels with age further supports heightened humoral immunity, potentially driven by chronic low-grade inflammation (inflammaging) and the accumulation of age-associated B cells (ABCs) [20, 21, 29–31]. However, the weak correlation between ANA and anti-ENA5 or anti-chromatin responses suggests that ANA positivity reflects a broader dysregulation of immune tolerance rather than a specific autoantigen-driven process.

In contrast to spontaneous autoimmunity, HgIA exhibited a striking age-related decline, accompanied by reduced immunoglobulin production and germinal center formation in old age mice, particularly in ANoA-negative individuals. This suggests that immunosenescence, characterized by a decline in T-cell-dependent B-cell responses [22–26], impairs the ability to mount a robust xenobiotic-induced humoral immune response. Notably, ANoA intensity in positive mice remained comparable across age groups, indicating that the genetic susceptibility to HgIA (e.g., MHC class II restriction) persists, but the threshold for initiating or sustaining this response increases in old age, likely due to diminished immune vigor. The large sample size was instrumental in uncovering this heterogeneity, revealing that even within an inbred strain, aging introduces variability in immune responsiveness to xenobiotics.

Flow cytometry analyses revealed significant age-related changes in T-cell subsets, with a decline in T cells and an increase in double-negative and double-positive T cells, consistent with age-related immune remodeling [60]. Mercury exposure augmented these changes in younger mice, promoting the differentiation of central to effector memory CD4^+^ T cells, mimicking age-related T-cell differentiation. In old age mice, however, these mercury-induced changes were blunted, reflecting the limited responsiveness of an aged immune system. The reduced frequency of CD4^+^ T cells in middle aged and old mice, coupled with high inter-individual variability, further underscores the impact of immunosenescence on xenobiotic-driven immunity.

The differential response to mercury in old age raises critical questions about the mechanisms underlying the failure to induce HgIA in a subset of animals. The association between ANoA positivity and higher IgG1 levels in old-age mice suggests that successful HgIA induction may depend on the preservation of T-cell-dependent B-cell function, specifically the maintenance of T_H_2 responses. Conversely, ANoA-negative mice exhibited reduced anti-chromatin antibodies, indicating that failure to mount HgIA may reflect a broader suppression of humoral immunity. This dichotomy could be linked to variability in the senescence-associated secretory phenotype (SASP) or the functionality of ABCs [29–31]. The observed heterogeneity in an inbred strain suggests that in genetically diverse populations, such as humans or outbred mouse models, the variability in immune responses to xenobiotics would be exponentially greater, necessitating further investigation into the interplay of genetic diversity and environmental exposures.

The clinical implications of these findings are profound, particularly for environmental exposures across the lifespan. While prolonged occupational exposure to xenobiotics like mercury is a known risk factor for autoimmunity [11], our data suggest that older individuals may be less susceptible to xenobiotic-induced autoimmune responses due to immunosenescence. However, the increased baseline autoantibody production in aging could exacerbate subclinical autoimmunity, potentially amplifying the effects of environmental triggers in susceptible individuals. This duality underscores the need for personalized approaches to assessing environmental risks in aging populations, particularly those with autoimmune predispositions.

Limitations of this study include the focus on a single xenobiotic (HgCl_2_) and a single mouse strain (B10.S), which may not fully capture the diversity of environmental exposures or genetic backgrounds relevant to human autoimmunity. The use of additional xenobiotics, such as silica or diesel exhaust particles [8–10], and diverse mouse models or human cohorts could further elucidate the generalizability of these findings. Additionally, examining cellular changes occurring in the peripheral blood would help characterize the systemic changes induced by HgCl_2_ exposure and improve the translational value of this study, Moreover, longitudinal studies tracking immune responses to repeated xenobiotic exposures across the lifespan could clarify the cumulative impact of environmental factors on age-related autoimmunity.

In conclusion, this study leverages a large sample size to reveal novel insights into the differential effects of aging on spontaneous and xenobiotic-induced autoimmunity. While aging promotes spontaneous autoantibody production, it dampens mercury-induced autoimmunity, likely due to immunosenescence. The observed heterogeneity in immune responses among inbred old mice suggests that genetic diversity and varied xenobiotic exposures would amplify this variability, highlighting the need for further research into the cellular and molecular mechanisms driving these responses. These findings have significant implications for understanding and managing autoimmune diseases in aging populations exposed to environmental triggers.

## Acknowledgements

We thank Joseph M. Christy for technical assistance in the early phases of this work. Research was supported by National Institutes of Health grants UH3ES027679 to K.M.P. This is publication #XXXXX from The Scripps Research Institute.

