## Supplemental Figure 1 for "Effect of Age on Xenobiotic-Induced Autoimmunity"

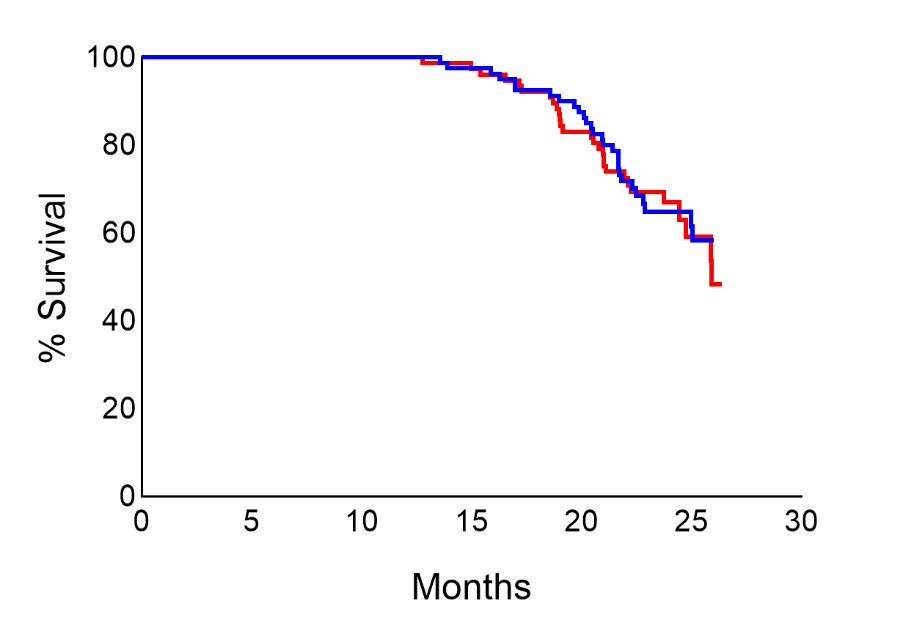

**Supplemental Figure 1.** **Survival curve for old B10.S mice.** 75% survival was reached at 21 months for female (red) (n=103) B10.S mice and 22 months for male (blue) (n=110) B10.S mice. Data is representative of three separate cohorts pooled together. Survival curves were created using the Kaplan and Meier method in Prism.
