## Supplemental Figure 2 for "Effect of Age on Xenobiotic-Induced Autoimmunity"

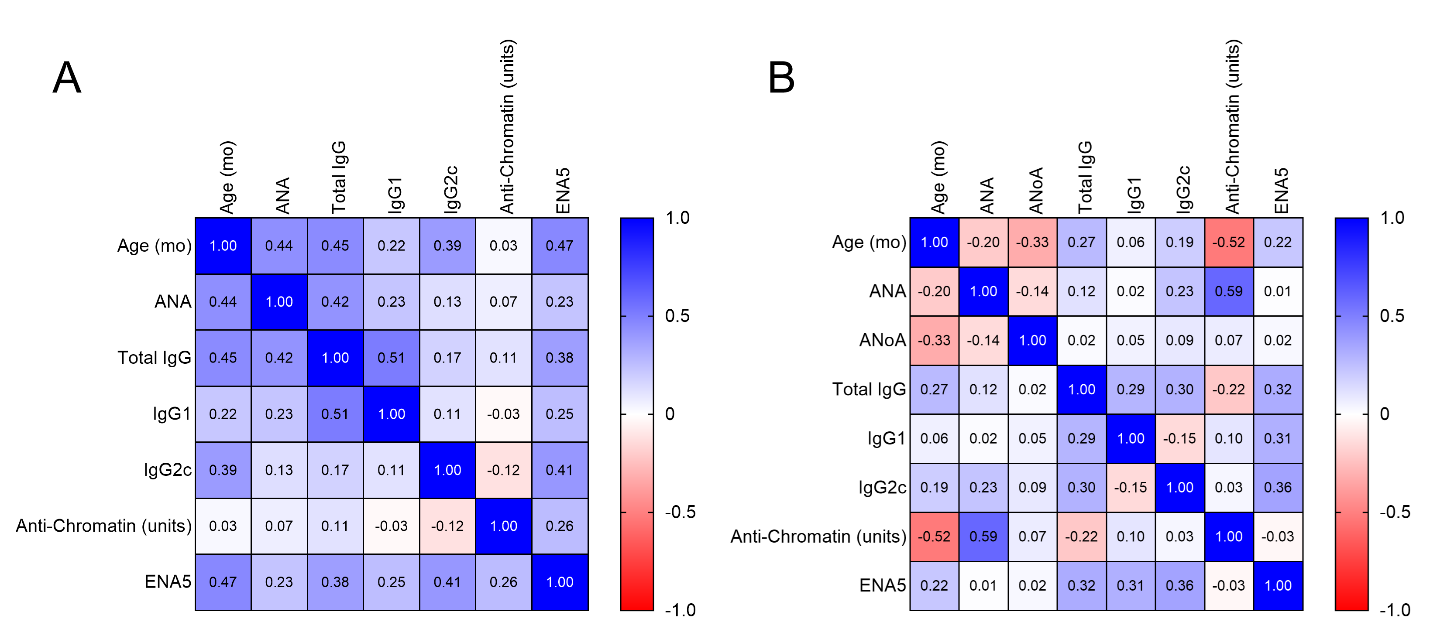
**Supplemental Figure 2. Correlation matrix of autoantibodies and immunoglobulins in B10.S mice.** A. mice treated with PBS, B. mice treated with HgCl_2_. Cell values represent Spearman’s rank correlation coefficient (r).
