## Supplemental Figure 3A for "Effect of Age on Xenobiotic-Induced Autoimmunity"

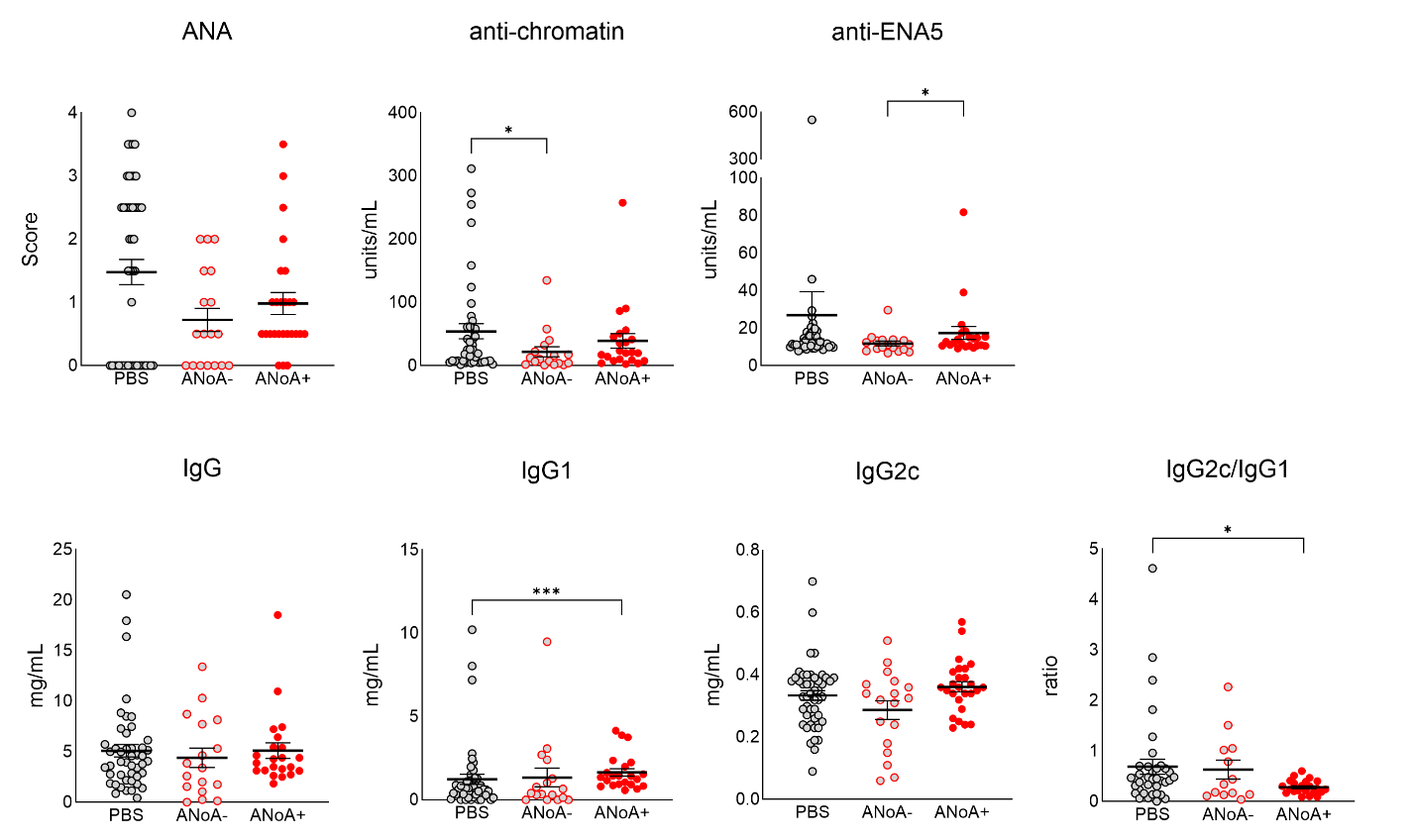
**Supplemental Figure 3A. Comparison of humoral immune responses in old PBS treated, ANoA negative and positive mice.** Sera assessment of 24 month old B10.S mice treated with PBS (n=49) and ANoA negative (n=18) and positive (n=26) mice treated with HgCl_2_. Statistical comparisons were performed with Mann-Whitney U tests. *P<0.05, **p<0.01, ***p<0.001, ****p<0.0001.
