## Supplemental Figure 3B for "Effect of Age on Xenobiotic-Induced Autoimmunity"

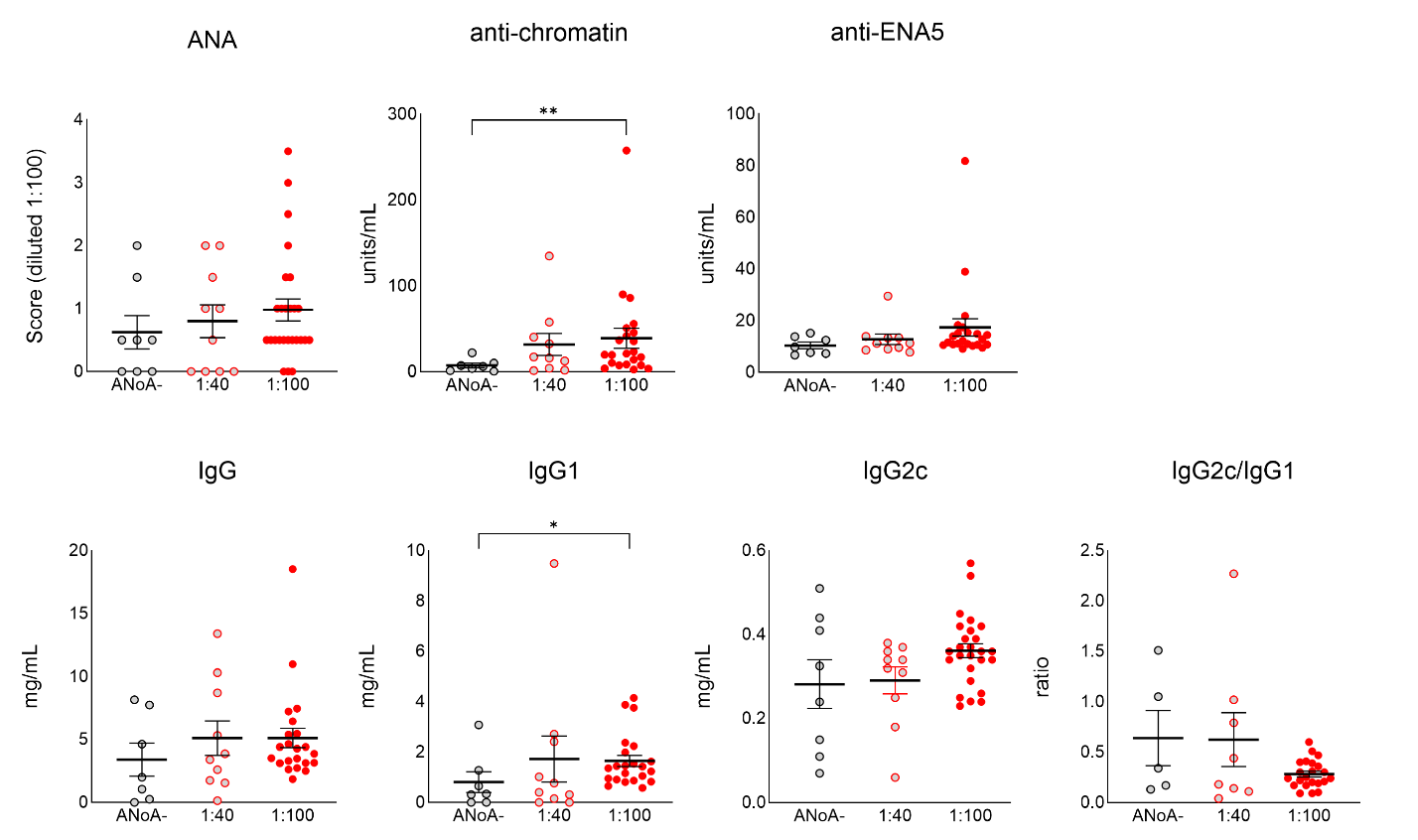
**Supplemental Figure 3B. Comparison of humoral immune responses in old HgCl_2_ treated mice that were ANoA negative and positive at 1:40 or 1:100 dilutions.** Sera assessment of 24 month old B10.S mice treated with HgCl_2_ that were ANoA negative at a 1:40 dilution (n=8), positive at a 1:40 dilution (n=10), and positive at a 1:100 dilution (n=26). Statistical comparisons were performed with Mann-Whitney U tests. *P<0.05, **p<0.01, ***p<0.001, ****p<0.0001.
