## Supplemental Figure 4 for "Effect of Age on Xenobiotic-Induced Autoimmunity"

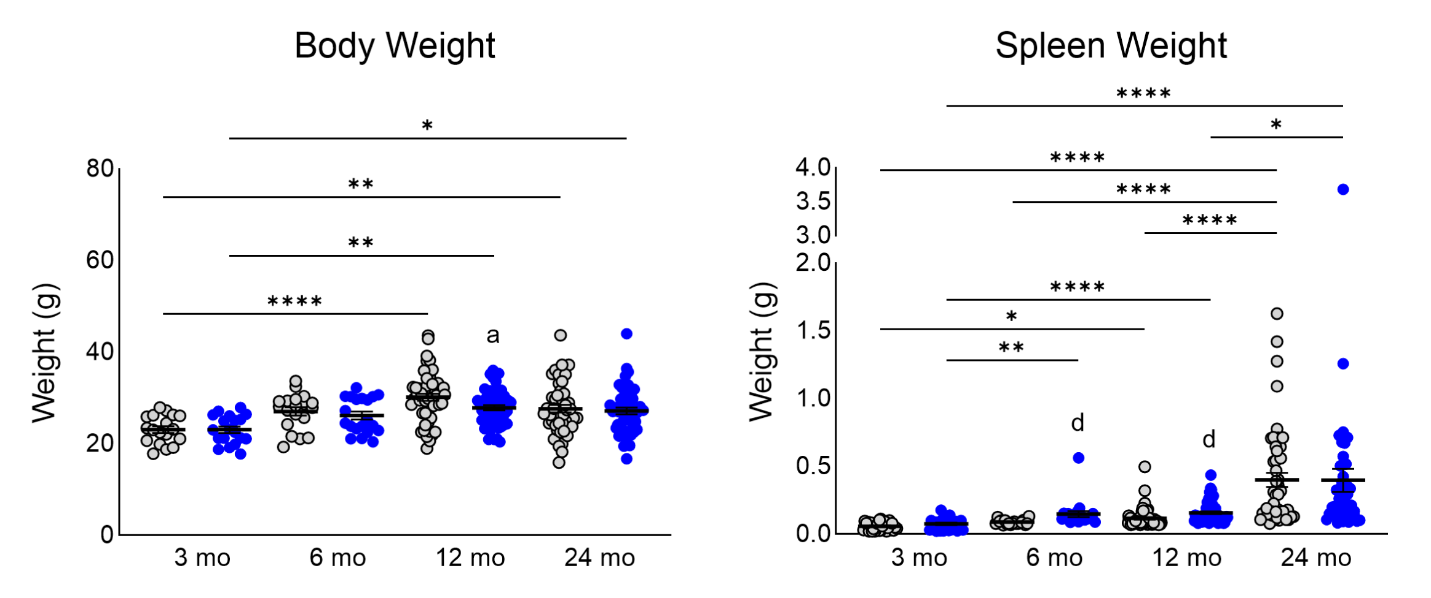
**Supplemental Figure 4. Age and HgCl_2_-induced changes in body and spleen weights in B10.S mice.** Measurements from B10.S mice at 3 (n=17/18), 6 (n=19/20), 12 (n=56/57) or 24 (n=48/44) months of age treated with PBS (gray) or HgCl_2_ (blue). Mice treatment and assays performed as described in Materials & Methods. Statistical comparisons between age groups were performed with Kruskal-Wallis tests. *P<0.05, **p<0.01, ***p<0.001, ****p<0.0001. Statistical comparisons of PBS and HgCl_2_ treatment groups were performed with Mann-Whitney U tests. a, p<0.05; b, p<0.01; c, p<0.001; d, p<0.0001.
