## Supplemental Table 1A for "Effect of Age on Xenobiotic-Induced Autoimmunity"

| **Marker** | **Conjugated flurophore** | **Clone** | **Manufacturer** |
| --- | --- | --- | --- |
| B220 | Pacific Blue | RA3-6B2 | BioLegend |
| CD19 | BUV 395 | 1D3 | BD Biosciences |
| CD93 | SB 436 | 'AA4.1 | Invitrogen |
| CD21/CD35 | PE/Cy 7 | 7E9 | BioLegend |
| CD23 | APC | B3B4 | BioLegend |
| IgD | BV 650 | 11-26c.2a | BioLegend |
| CD38 | PE/Dazzle 594 | 90 | BioLegend |
| CD86 | BV 785 | GL-1 | BioLegend |
| CD80 | PE/Cy 5 | 16-10A1 | BioLegend |
| CD69 | BV 711 | H1.2F3 | BioLegend |
| GL7 | AF 488 | GL7 | BioLegend |
| CD138 | BV 421 | 281-2 | BioLegend |
| TCRb | AF 647 | H57-597 | BioLegend |
| CD4 | BV 750 | GK1.5 | BioLegend |
| CD8a | PerCP/Cy 5.5 | 53-6.7 | BioLegend |
| CD44 | BV 480 | IM7 | BD Biosciences |
| CD62L | APC/Fire 750 | MEL-14 | BioLegend |
| CD11b | BV 570 | M1/70 | BioLegend |
| CD11c | BB 515  Spark NIR 685 | N418 | BD Biosciences  BioLegend |
| IgM | BUV 661 | 5153-238 | BD Biosciences |
| T-bet | PE | 4B10 | BioLegend |
| Viability | DAPI  Ghost Dye UV 450 |  | Cytek |

**Supplementary Table 1A. List of conjugated antibodies used in flow cytometry.** Panel was updated for follow-up study of age-associated B cells, changes are in red.
