## Supplemental Table 1B for "Effect of Age on Xenobiotic-Induced Autoimmunity"

| **Marker** | **Conjugated flurophore** | **Clone** | **Manufacturer** |
| --- | --- | --- | --- |
| CD19 | BUV 395 | 1D3 | BD Biosciences |
| CD38 | PE/Dazzle 594 | 90 | BioLegend |
| CD86 | BV 785 | GL-1 | BioLegend |
| CD80 | PE/Cy 5 | 16-10A1 | BioLegend |
| CD69 | BV 711 | H1.2F3 | BioLegend |
| GL7 | AF 488 | GL7 | BioLegend |
| TCRb | AF 647 | H57-597 | BioLegend |
| CD4 | BV 750 | GK1.5 | BioLegend |
| CD8a | PerCP/Cy 5.5 | 53-6.7 | BioLegend |
| CD44 | BV 480 | IM7 | BD Biosciences |
| CD62L | APC/Fire 750 | MEL-14 | BioLegend |
| CD11b | BV 570 | M1/70 | BioLegend |
| ICOS | PE/Cy 7 | 7E.17G9 | BioLegend |
| PD-1 | PE/Fire 700 | 29F.1A12 | BioLegend |
| CD49b | RB 744 | HMa2 | BD Biosciences |
| CXCR5 | PE | L138D7 | BioLegend |
| Viability | eFluor 780 | - | Invitrogen |

**Supplemental Table 1B. List of conjugated antibodies used in flow cytometry.** Panel used for the follow-up study of germinal center T follicular helper cells.
