## Supplemental Table 2 for "Effect of Age on Xenobiotic-Induced Autoimmunity"

| Cell type | Phenotype |
| --- | --- |
| T | TCRb+ |
| CD4 T | TCRb+ CD4+ CD8a- |
| CD8 T | TCRb+ CD4- CD8a+ |
| Naïve CD4 T | TCRb+ CD4+ CD8a- CD44- CD62L+ |
| Central Memory CD4 T | TCRb+ CD4+ CD8a- CD44+ CD62L+ |
| Effector Memory CD4 T | TCRb+ CD4+ CD8a- CD44+ CD62L- |
| Naïve CD8 T | TCRb+ CD4- CD8a+ CD44- CD62L+ |
| Central Memory CD8 T | TCRb+ CD4- CD8a+ CD44+ CD62L+ |
| Effector Memory CD8 T | TCRb+ CD4- CD8a+ CD44+ CD62L- |
| B | CD19+ |
| B1 | CD19+ CD93- B220- |
| Transitional B | CD19+ CD93+ B220lo |
| B2 | CD19+ CD93- B220+ |
| Follicular B | CD19+ CD93- B220+ CD23+ CD21+ |
| Memory B | CD19+ CD93- B220+ CD23- CD21- |
| Class-switched Memory B | CD19+ CD93- B220+ IgD- CD21- |
| Marginal Zone B | CD19+ CD93- B220+ CD23- CD21hi |
| Germinal Center B | CD19+ CD93- B220+ CD23- CD21- GL7hi CD38lo |
| Plasma Cells | CD138+ B220- |
| Plasmablasts | CD138+ B220+ |
| nonTB | TCRb- CD19- |
| Myeloid | TCRb- CD19- CD11b+ CD11c- |
| Monocyte | TCRb- CD19- CD11b+ CD11c- SSClo |
| Granulocyte | TCRb- CD19- CD11b+ CD11c- SSChi |
| Dendritic | TCRb- CD19- CD11chi |
| DC1a | TCRb- CD19- CD11chi CD11b+ CD8a- |
| DC2 | TCRb- CD19- CD11chi CD11b- CD8a+ |

**Supplemental Table 2. Cell phenotype chart.** List of cell surface markers used to identify cell types.
