## Supplemental Table 3 for "Effect of Age on Xenobiotic-Induced Autoimmunity"

| ***Posture* – kyphosis, lordosis, scoliosis:** | |
| --- | --- |
| 0 | normal posture |
| 1 | slight alteration in posture |
| 2 | moderate alteration in posture |
| 3 | severe alteration in posture |
| 4 | severe alteration in posture which impairs animals ability to move about, groom, eat or drink |
| ***Mobility*:** | |
| 0 | normal activity |
| 1 | walks with only minimal change in gait |
| 2 | walks with stiff gait |
| 3 | walks with stilted gait, with limited |
| 4 | limited mobility/activity due to pronounced stiffness or loss of range of motion |
| ***Haircoat/Skin (trunk)*:** | |
| 0 | normal haircoat, skin |
| 1 | alopecia with no skin lesions, mild seborrhea |
| 2 | superficial dermatitis involving less than 10% of body |
| 3 | superficial dermatitis involving >10% but <30% of body |
| 4 | severe dermatitis, superficial dermatitis over 30% of body or full thickness skin lesions extending into subcutis and or muscle layers |
| ***Haircoat/Skin (head)*:** | |
| 0 | normal haircoat, skin, pinnae |
| 1 | alopecia with no skin lesions, mild seborrhea |
| 2 | superficial |
| 3 | mild superficial dermatitis involving muzzle or superficial dermatitis involving >5% and < 20% of face and/or pinnae |
| 4 | moderate superficial dermatitis involving muzzle, severe dermatitis involving >20% of face and/or ears or full thickness skin lesions extending into subcutis and or muscle layers |
| ***Haircoat/Skin (extremities/limbs)*:** | |
| 0 | normal haircoat, skin |
| 1 | alopecia with no skin lesions, mild seborrhea |
| 2 | superficial dermatitis involving < 5% of limb |
| 3 | superficial dermatitis >5% and < 20% of limb, mild swelling or mildly affected gait |
| 4 | moderate to severe superficial dermatitis which includes swelling and impairment of mobility, severe dermatitis involving >20% of limb or full thickness skin lesions extending into subcutis and or muscle layers |
| ***Ocular Lesions:*** | |
| 0 | normal appearance of eye, lids |
| 1 | alopecia without swelling or skin lesions, cataract formation |
| 2 | mild blepharitis involving one eye |
| 3 | mild blepharitis involving both eyes, moderate blepharitis of one eye, mild corneal edema with no other ocular lesions |
| 4 | moderate to severe blepharitis involving both eyes, any active corneal lesions (neovascularization, ulceration, pannus), any periorbital swelling consistent with tumor or abscessation, buphthalmous |
| ***Body Weight:*** | |
| 0 | normal body weight |
| 1 | weight change (increase/decrease) within 5% of previously recorded stable baseline weight |
| 2 | mild weight change (increase/decrease) within 10% of previously recorded stable baseline weight |
| 3 | moderate weight change (increased/decreased) within 15% of previously recorded stable baseline weight |
| 4 | severe weight change (increased/decreased) greater than 15% of previously recorded stable baseline weight |
| ***Other injury/trauma/impairment:*** | |
| 0 | normal |
| 1 | small healing wound |
| 2 | mild wounding due to aggression or other trauma |
| 3 | moderate wounding due to aggression, trauma |
| 4 | severe wounding due to aggression, trauma |
| ***Dehydration:*** Loss of skin turgor | |
| ***Anemia:*** Pale coloration | |
| ***Respiratory Distress:*** Cyanotic coloration; Labored respiration; Increased respiration | |
| ***Hypothermia:*** Piloerection; Shivering | |
| ***Other severe conditions:*** Paralysis, Rectal prolapse; Penile prolapse; Neurological signs: head tilt; tremors, seizures, ataxia | |
| **Euthanasia is indicated for the following conditions:** | |
| 1.            Any animal that exhibits any grade 4 clinical signs as listed above. | |
| 2.            Any animal that exhibits grade 3 clinical signs for which therapy is not provided or the animal is non-responsive to therapy. | |
| 3.            Any animal that exhibits a progression of signs to grade 3 for which therapy is not provided or the animal is non-responsive to therapy. | |
| 4.            Any animal exhibiting clinical signs listed in items “i” through “m” above which are non-responsive to therapy or for which therapy is unrewarding as in cases of neurological impairment, rectal prolapse, penile prolapse. | |
| 5.            Any animal which has met euthanasia criteria as outlined in other TSRI guidelines (e.g. tumor development or arthritis). | |

**Supplementary Table 3. Monitoring parameters for animals used in aging research.** List of monitoring parameters used to evaluate morbidity.
